# Shroom3-dependent pillar cell constriction establishes a mechanical threshold for respiratory lamellar stability

**DOI:** 10.64898/2026.07.17.739096

**Authors:** Mathieu Preußner, Alisa Kahler, Marion Basoglu, Anna Mertens, Daniela S. Krause, Stefan Eimer, Virginie Lecaudey

## Abstract

How developing organs acquire complex architectures while maintaining tissue integrity under mechanical stress is a fundamental question in mechanobiology. In the gill, pillar cells support the thin respiratory lamellae, but how they withstand hemodynamic forces is unknown.

Here, we show that actomyosin-rich contractile apparatuses (CA) in zebrafish pillar cells act as load-bearing units that generate contractile forces and stabilize the lamellae through adhesion to and enclosure of extracellular collagen columns. CA number correlates with pillar cell constriction, linking subcellular architecture to tissue geometry. We further identify Shroom3 and Vinculin b as key regulators of CA assembly and force transmission: *shroom3* deficiency or loss of *vclb* impairs CA formation and reduces pillar cell constriction, while loss of *shroom3* causes lamellar collapse.

Together, these findings support a mechanical threshold model in which lamellar integrity depends on a minimum number of CAs per pillar cell and establish the gill lamellae as a model for linking subcellular contractility to tissue-scale mechanical stability.

## Introduction

To form complex multicellular organs, cells must proliferate, change shape, adhere to one another and to the extracellular environment, and migrate. The spatiotemporal coordination of these processes is essential and is controlled by robust gene regulatory programs. At the same time, tissue architecture is not only genetically encoded but also mechanically regulated by intrinsic and extrinsic forces, including tissue stiffness, cell density, blood flow and hydrostatic pressure, which continuously influence organ formation and homeostasis ^1–3^.

At the core of cellular force generation and mechanotransduction is the cytoskeleton. In particular, the dynamic actomyosin network enables cells to generate, sense and transmit mechanical forces through coupling to neighboring cells and the extracellular matrix (ECM), thereby driving tissue morphogenesis and maintaining tissue architecture ^4,5^. A fundamental mechanism underlying morphogenesis is apical constriction, an evolutionary conserved cell shape change that occurs in many developmental contexts, including gastrulation and neural tube closure ^4,6,7^. In epithelial cells, apically localized actomyosin generates contractile forces that transform columnar cells into wedge-shaped cells. When coordinated across groups of cells, this process can drive the cellular rosette formation, tissue bending and epithelial invagination ^8–11^.

Shroom3 is a key regulator of epithelial apical constriction in vertebrates. First identified as the gene responsible for neural tube closure defects in mouse *Mushroom* mutants ^7^, Shroom3 was subsequently shown to be required for constriction-based morphogenesis in several developmental contexts, including gut formation, lens placode invagination, and neuromast formation ^12–14^. Beyond development, SHROOM3 has also been implicated in pathological processes, including chronic kidney disease, as it is required to maintain the glomerular filtration barrier ^15–18^. Shroom proteins are characterized by conserved domains, including an N-terminal PDZ domain and two Apx/Shrm domains (ASD), which mediate interactions with actin and Rho-associated kinase (ROCK), respectively. Mechanistically, Shroom3 localizes to the apical side of epithelial cells through its interaction with actin and recruits ROCK1/2 proteins, which in turn phosphorylate the myosin regulatory light chain of non-muscle myosin II (NMII) to initiate apical constriction ^19,20^.

We previously showed that *shroom3* is required for the formation of radial rosettes that prefigure sensory neuromasts in the posterior lateral line (pLL) ^14^. Although *shroom3* homozygous mutants survive to adulthood, they exhibit a conspicuous gill phenotype, in which the opercula appear more open, suggesting an underlying defect in gill architecture.

During respiration, the operculum opens as the mouth closes, creating a pressure drop in the branchial cavity that drives water flow across the gills. Each gill contains four arches on each side, each bearing two alternating rows of filaments, medial and lateral, that extend outward from the arch (Fig. 1A-B).

**Fig. 1.**
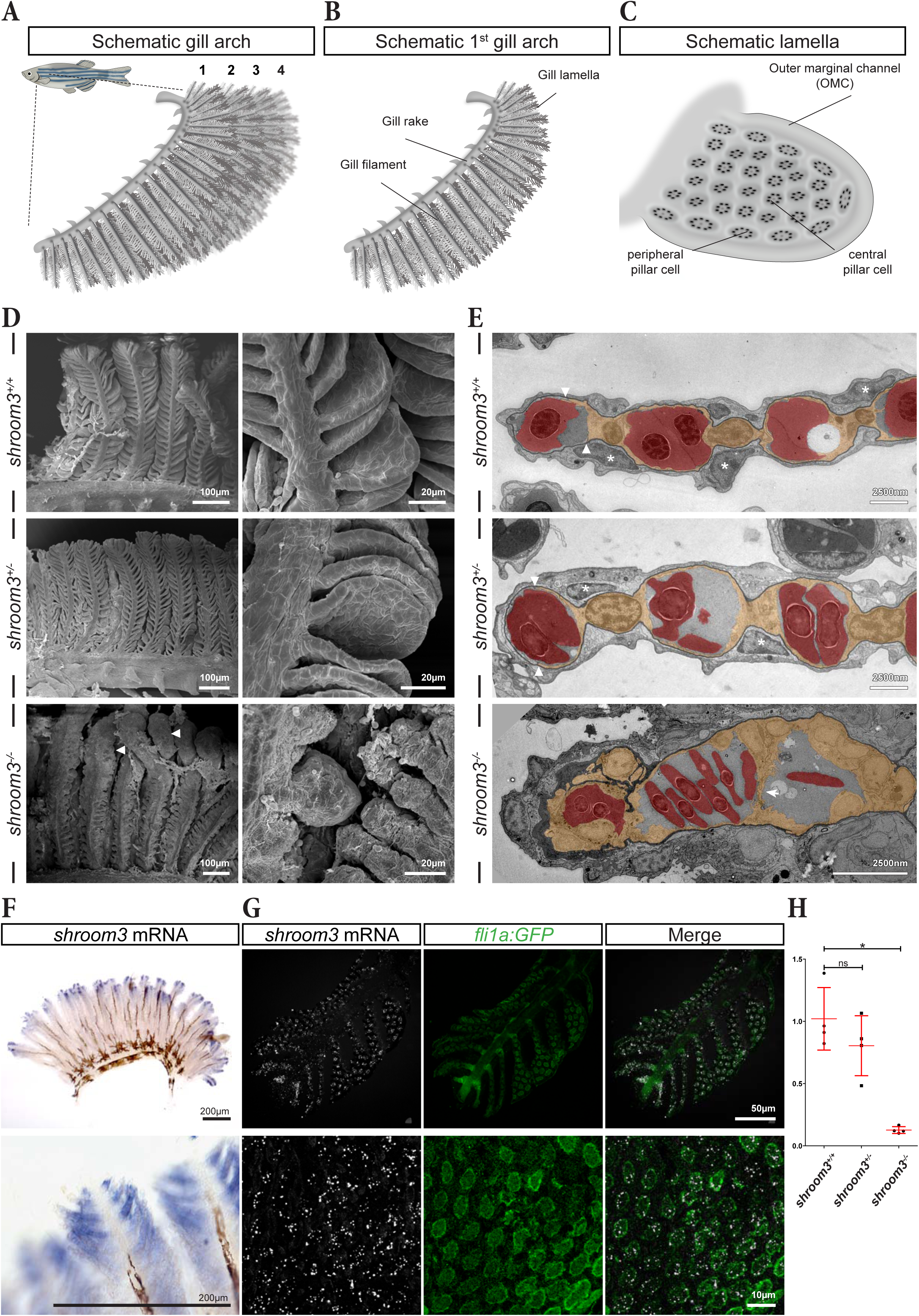
*shroom3* is expressed in pillar cells and is required for lamellar morphogenesis in the gills. (A) Scheme of the four gill arches in the adult zebrafish. (B) Schematic of a single gill arch. (C) Schematic of a single lamella. (D) Scanning electron micrographs of a gill arch region containing several filaments (left) and higher-magnification views of lamellae from the indicated *shroom3* genotypes. (E) Transmission electron micrographs of single lamellae in cross-section from the indicated *shroom3* genotypes. Erythrocytes, pillar cells (PCs) and the pavement cell (PVC) basement membrane are pseudocolored in red, yellow and black, respectively. (F) Colorimetric *in situ* hybridization for *shroom3* on a whole gill arch (top) and higher-magnification view of a filament showing that *shroom3* expression is restricted to lamellae at the distal tip. (G) Gill cryosections from a *fli1a:GFP* fish stained for *shroom3* mRNA by HCR (top panels), with higher-magnification views showing *shroom3* expression specifically in GFP-positive PCs within the lamellae. Shown are maximum-intensity projections (MIPs) of confocal stacks. (H) RT-qPCR analysis of dissected adult gills showing relative *shroom3* expression across the indicated *shroom3* genotypes. Error bars indicate mean ± SD. Two-tailed Mann-Whitney U test: \**P* ≤ 0.05.

Along each filament, lamellae branch out bilaterally in alternating rows and serve as the main sites of gas exchange (Fig. 1B-C). These lamellae are continuously exposed to internal hemodynamic and external water pressure, and their integrity therefore depends on resident cells capable of stabilizing the tissue under mechanical load.

Pillar cells (PCs) form a highly organized vascular system and are strategically positioned between the opposing epithelial layers of pavement cells (PVCs) that cover the lamellae. PCs consist of a central cylindrical cell body from which flat cytoplasmic arms extend from both ends and connect with neighboring PCs, therefore forming a continuous cellular network that encloses a continuous vascular space (VS) within each lamella ^21^. A defining feature of PCs is the presence of collagen columns (CCs) arranged in a ring around the nucleus, and oriented perpendicular to the epithelial PVCs. CC are continuous with the PVC basement membrane and are enfolded by the PC plasma membrane. On the cytoplasmic side, each CC is surrounded by actomyosin-rich cables called contractile apparatuses (CAs) ^21–23^. In zebrafish, PCs were first described morphologically in the 80s ^24^, and were more recently shown to derive from cranial neural crest cells ^25,26^. These unique characteristics have led to the hypothesis that PCs act as active mechanical elements within the lamellae, generating contractile forces through their actomyosin-rich CAs to redistribute lamellar blood flow and regulate gas exchange ^21,22^.

Here we provide a molecular and quantitative characterization of PCs in *Danio rerio* and identify their spatial organization as a key determinant of lamellar architecture. We show that Shroom3 is required for PCs morphogenesis and CA assembly, and that both Shroom3 and Vinculin b are essential for maintaining lamellar stability. Our data support a threshold model in which lamellar integrity depends on the number and organization of CAs within each PC. Below a critical threshold, the forces generated and transmitted by the CA-CC system are insufficient to withstand hemodynamic pressure, resulting in lamellar collapse and ballooning. Together, these findings link Shroom3-dependant contractility and Vinculin b-mediated force transmission to tissue-scale stability in a physiologically dynamic organ.

## Results

### *shroom3* is expressed in pillar cells and is required for lamellar morphogenesis in the gills

To characterize the gill phenotype of *shroom3* mutants, we first examined gill ultrastructure by scanning electron microscopy (SEM). SEM confirmed the overall organization of the gills, with multiple filaments extending from each gill arch and bearing regularly spaced lamellae arranged in a staggered pattern on both sides (Fig. 1A-D). In wild-type specimens (*shroom3^+/+^*) the PVCs covering the lamellar surface were readily visible (Fig. 1D, right panel). No obvious structural differences were observed between *shroom3^+/+^* and *shroom3^+/-^*gills. In contrast, lamellae in *shroom3^-/-^* mutants were severely affected, exhibiting a swollen, ballooned morphology, with apparent fusion between neighboring lamellae, particularly at the filament tip (arrowheads in Fig. 1D).

To further analyze the cellular organization, we performed transmission electron microscopy (TEM) on lamellae (Fig. 1E). In *shroom3^+/+^* fish, PCs (yellow in Fig. 1E) were regularly spaced and exhibited their characteristic morphology, including thin cytoplasmic arms extending from the cell body to connect adjacent PCs. This arrangement defined the vascular spaces (VSs) ^24^, which were filled with nucleated erythrocytes (red in Fig. 1E). At the distal end of the lamellae, the cytoplasmic arms of the most distal PC made direct contact with an endothelial cell (EC) (arrowhead in Fig. 1E) thereby forming the outer marginal channel (OMC). The entire lamellae were covered by PVCs, whose large, flattened nuclei were predominantly positioned above the PC nuclei (asterisks in Fig. 1E). *shroom3^+/-^* lamellae showed an overall similar cellular arrangement although they appeared slightly wider, with enlarged PC cell bodies and VSs. In contrast, lamellae in *shroom3^-/-^* fish were severely disrupted. The regular arrangement and alternating pattern of PCs and VSs were lost. Consequently, the vascular compartment was markedly expanded and disorganized, occupying the central core of the lamella (Fig. 1E), in which erythrocytes accumulated. PCs were located at the periphery of the lamella surrounding the enlarged VS, maintaining only limited residual contact (arrow in Fig. 1E). Additionally, the basement membrane (BM) appeared thickened (black in Fig. 1E) and PVCs were overcrowded (Fig. 1E).

We next asked where *shroom3* is expressed in the gills. Colorimetric *in situ* hybridization on dissected gill arches revealed strong expression at the distal tip of filaments (Fig. 1F). However, the resolution was insufficient to determine whether the signal was specific to PCs or was also present in the surrounding epithelium (Fig. 1F, lower panel). To overcome this limitation, we performed fluorescent *in situ* hybridization chain reaction (HCR). This more sensitive approach demonstrated that *shroom3* was expressed in PCs, as indicated by overlap with the PC marker *fli1a:GFP* (Fig. 1G) ^27^. Furthermore, *shroom3* expression was detected in lamellae along the entire filament (Fig. 1G, top panels), suggesting that the distal restriction of the signal observed with colorimetric ISH most likely reflected limited probe penetration. RT-qPCR performed on dissected gills confirmed *shroom3* expression and revealed a strong reduction in transcript levels in mutants, consistent with nonsense-mediated mRNA decay (Fig. 1H).

Together, these data indicate that *shroom3* is expressed in lamellar PCs and is required for their structural organization and cellular arrangement, supporting a key role for Shroom3 in maintaining the organization of the gill respiratory surface.

### Pillar cells form actomyosin-rich contractile apparatuses linked to collagen columns

Having established that *shroom3* is expressed in PCs, we next set out to characterize PC morphology in wild-type fish using markers of the actomyosin cytoskeleton, cell junctions and cell-matrix adhesions, thereby establishing a reference for comparison with *shroom3* mutants. For this, we drew on previous work in cultured pufferfish (*Takifugu rubripes)*, another actinopterygian species ^21^.

Phalloidin staining revealed dense actin rings surrounding each PC nucleus (Fig. 2). At higher resolution, these rings were found to consist of 6 to 8 smaller, closely packed circular structures resembling a string of beads (Fig. 2A). Staining with fluorescent Concanavalin A, previously shown to label CCs ^23^, revealed a small dot at the centre of each bead, indicating the presence of CCs surrounded by the actin-rich CA of PCs (Fig. 2A, close-up). Consistent with this interpretation, in the absence of CC staining, the CA appeared as a hollow actin structure with a central unstained space (Fig. S1A). Orthogonal projections of confocal stacks further revealed that both the CAs and the CCs extend perpendicular to the lamellar surface along the full height of the PCs (Fig. 2A, longitudinal view). Due to the close apposition of the CA, the PC nuclei exhibited corresponding indentations, giving them a lobed, oak leaf-like morphology (Fig. S1A).

**Fig. 2.**
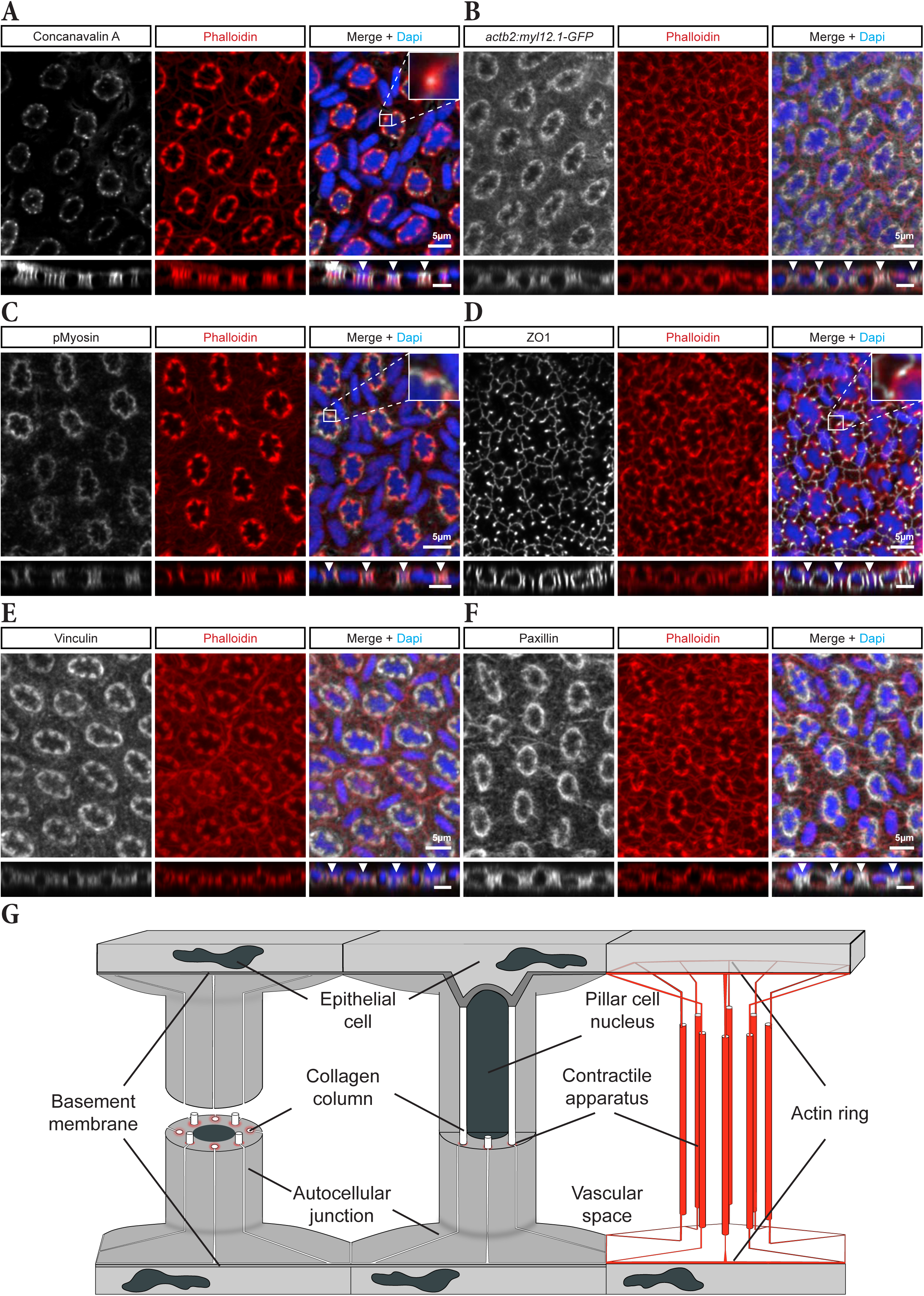
Pillar cells form actomyosin-rich contractile apparatuses linked to collagen columns. (A-F) MIPs of cryosections from adult gill arches showing the molecular organization of PCs within the lamellae. For each marker, the left panel shows the staining, transgene or immunolabeling as indicated, the middle panel shows the contractile apparatus (CA) visualized with phalloidin, and the right panel shows the merged image including DAPI-stained nuclei. Below each panel is an orthogonal projection of the confocal stack for the respective markers (3-µm MIP). Individual PCs are marked by arrowheads. Markers analyzed were Concanavalin A (A), *myl12.1-GFP* (B), phospho-myosin light chain (pMLC) (C), ZO-1 (D), Vinculin (E), Paxillin (F). (G) Schematic representation of PC architecture, highlighting the actin-rich CA in red.

Non-muscle myosin (NMII) visualized using the *Tg(actb2:myl12.1-EGFP)* transgene (hereafter *myl12-GFP*), co-localized with actin at the CA (Fig. 2B). Immunostaining for phosphorylated myosin further marked the CA, consistent with an active contractile state (Fig. 2C). Notably, the hollow actin structure corresponding to the CA, were bordered on the outer side, opposite the nucleus, by tight junctions (TJs), as shown by Zonula occludens-1 (ZO-1) immunostaining (Fig. 2D, inset, S1C-E). This is consistent with a role for TJs in sealing the CA around the CC at autocellular junctions ^21^.

ZO-1 staining further revealed a prominent TJ network connecting all autocellular junctions of a given PC to an outer junctional ring encircling the cell (Fig. 2D). This organization indicates that TJs form a continuous network that seals the vascular space, likely preventing blood leakage. Consistent with this architecture, phalloidin staining also revealed an outer actin ring that mirrored the TJ network (actin ring in Fig. 2D,G) and connected the CAs to cell-cell junctions between PCs (Fig. 2D,G).

To further investigate cytoskeletal organization, we examined proteins involved in cell–ECM adhesion. Immunostaining for the focal adhesion components Vinculin and Paxillin, two key adaptor proteins linking Integrins to the actin cytoskeleton, revealed their localization at the CA surrounding the CCs (Fig. 2E,F). In addition, GFP signal was detected at CCs in the endogenous knock-in line *TgKI(itgb1b:itgb1b-sGFP)* (hereafter *itgb1b-sGFP*, Fig. S1B) ^28^. Together, these observations suggest that Integrin β1b-containing adhesion complexes at the PC surface mechanically couple the extracellular CC to the intracellular CA via adaptor proteins including Paxillin and Vinculin (schematized in Fig.S1C-E).

### Shroom3 is required for contractile apparatus formation and pillar cell constriction

To investigate the basis of lamellar disorganization in *shroom3* mutants, we next examined PC morphology across genotypes. Combined Myl12-GFP and Phalloidin staining revealed no obvious qualitative defects in CA organization in shroom3^+/-^ PCs compared with wild-type controls (Fig. 3A). In contrast, homozygous mutants exhibited complete disruption of the CA network, with no structures resembling the wild-type cellular architecture. Instead, the actomyosin scaffold was displaced toward the periphery of the lamella, suggesting a failure of CA assembly and/or maintenance (Fig. 3A). As a consequence, the regularly organized vascular network collapsed into a single enlarged vascular space filled with blood cells, consistent with the defects observed by TEM (Fig. 1E).

**Fig. 3.**
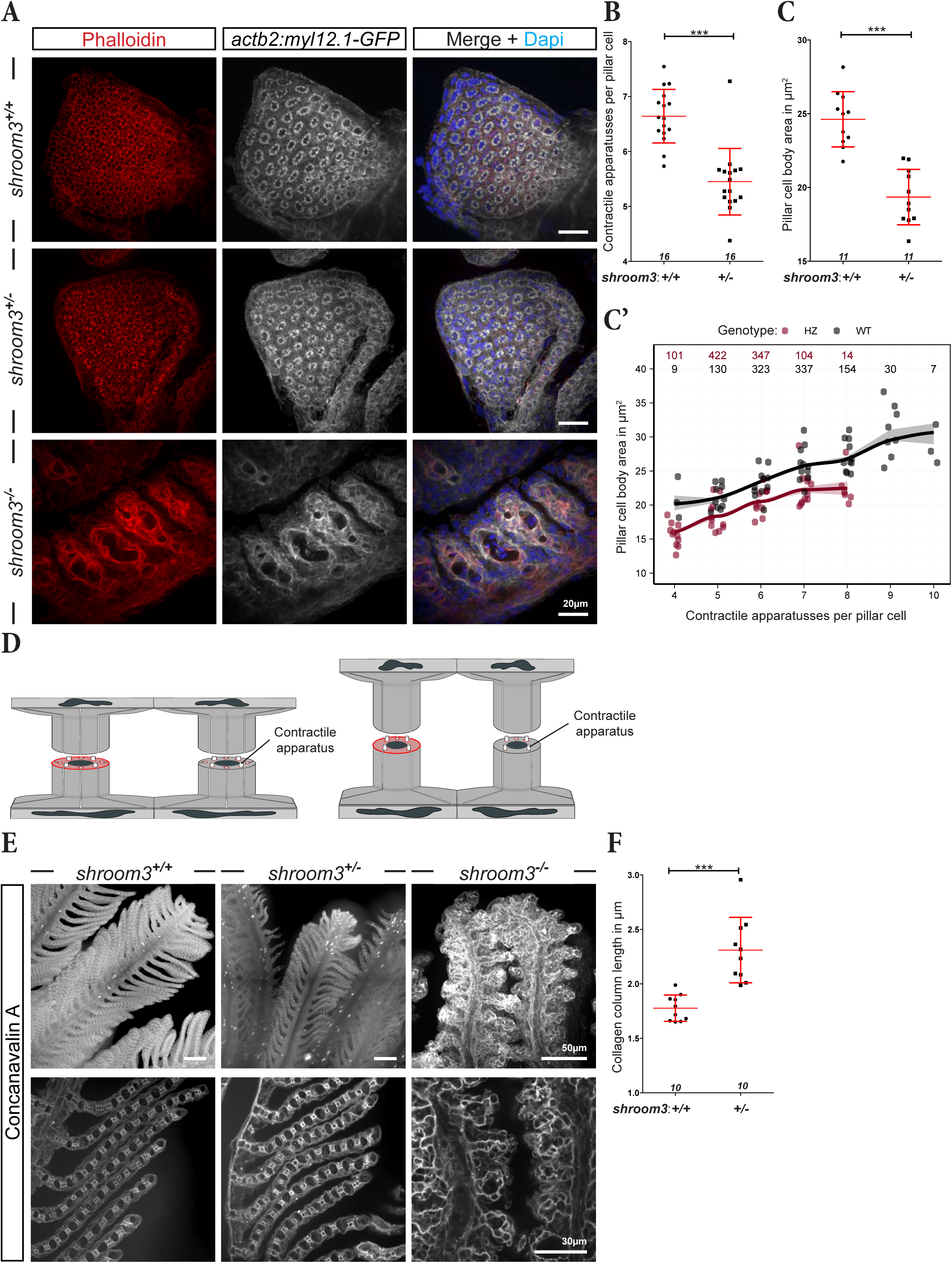
Shroom3 is required for contractile apparatus formation and pillar cell constriction. (A) Representative MIPs of cryosections from adult gill arches expressing *myl12.1-GFP*, labeled with phalloidin (F-actin) and DAPI (nuclei), highlighting the actomyosin-rich contractile apparatus of PCs across all *shroom3* genotypes, as indicated. (B) Quantification of CA number per PC in *shroom3 ^+/+^* and *shroom3^+/-^* fish. (C) Quantification of PC cross-sectional body area in *shroom3 ^+/+^* and *shroom3 ^+/-^* fish. (C’) PC body area plotted as a function of CA number per PC. Each point represents the mean body area of PCs with a given CA count within one animal, color-coded by genotype. Colored lines show genotype-specific LOESS-fitted mean values, with shaded ribbons indicating SEM. Numbers below the x axis indicate CA number. Numbers at the top indicate the number of PCs analyzed in each CA category for each genotype. (D) Schematic illustrating measurement of PC cross-sectional body area (red area) and interpretation of this readout. More constricted PCs exhibit a larger cell body area in cross-section and, on average, more CAs (left), whereas less constricted PCs display a smaller area (right) and, on average, fewer CAs. (E) Representative MIPs of confocal stack from ECi-cleared gills stained with Concanavalin A to label collagen columns (CCs) in the indicated *shroom3* genotypes (top). MIPs at higher magnification, showing individual PCs within lamellar cross-sections, are shown below. (F) Quantification of CC length in *shroom3^+/+^* and *shroom3 ^+/-^* fish. (B, C, F) Error bars indicate mean ± SD. Each data point represents the mean value of 90 PCs from three images of one fish. Numbers at the bottom of each panel indicate the number of animals analyzed (n). Two-tailed Mann-Whitney U-test : \*\*\**P* ≤ 0.001, ns = not significant.

Because the severe *shroom3^-/-^* phenotype precluded detailed quantitative analysis of lamellar architecture, subsequent morphometric analyses focused on heterozygous animals, in which overall lamellar organization was retained. Quantification of Phalloidin-stained cryosections revealed that *shroom3^+/-^* PCs contained, on average, one fewer CA per cell than wild-type siblings (Fig. 3A,B).

To determine whether this reduction in CA number affects PC constriction, we quantified PC body area, defined as the cross-sectional area enclosed by the CA in individual cells (Fig. 3C,D). Assuming constant cell volume, reduced constriction along the PC long axis, parallel to the CA, is expected to result in increased PC height and decreased cell body area, as previously reported ^29^(Fig. 3D). Consistent with reduced longitudinal constriction, *shroom3^+/-^* PCs displayed smaller body areas than wild-type siblings (Fig. 3C).

We next asked whether this phenotype could be explained solely by the reduction in CA number, or whether Shroom3 also contributes more directly to CA contractile function. To distinguish between these possibilities, we compared PCs with the same number of CAs across genotypes (Fig. 3C’). For a given CA number, *shroom3^+/+^* PCs consistently exhibited larger cell body areas than *shroom3^+/-^*PCs, suggesting that Shroom3 promotes PC constriction not only by regulating CA number, but also by contributing to the contractile state of the CA. In both genotypes, CA number positively correlated with PC body area, with cells containing more CAs exhibiting larger areas than those with fewer CAs (Fig. 3C’). These data suggest that the number of CAs per PC is an important determinant of PC constriction. Notably, the variation in CA number was reflected spatially in both genotypes, with PCs in the centre of the lamellae containing fewer CAs than PCs at the periphery (Fig. S2A-C). Interestingly, both central and peripheral PCs contained fewer CAs in *shroom3^+/-^* lamellae compared to wild-type siblings (compare Fig. S2B to S2C).

To independently and more directly assess the constriction state of PCs, we additionally measured the length of the CCs labelled with Concanavalin A in cleared adult gill arches (Fig. 3E,F). As an internal control, we first compared central and peripheral PCs, reasoning that if CA number predicts constriction state, peripheral PCs should be more constricted. Consistent with this idea, peripheral PCs had significantly shorter CCs than central PCs, supporting CC length as an independent and more direct readout of PC constriction (Fig. S2D,E). In *shroom3^+/-^* mutants, CCs were significantly longer than in wild-type siblings (Fig. 3E,F), confirming reduced PC constriction. This finding was also consistent with the increased lamellar height observed by TEM in *shroom3^+/-^* mutants (Fig. 1E; Fig. S3). In *shroom3^-/-^* mutants, CCs were not detectable (Fig. 3E), consistent with the absence of recognizable CA (Fig. 3A) and the severe disruption of lamellar architecture observed by TEM. Nonetheless, small extracellular accumulations, presumably remnants of CCs, could still be observed (arrowheads in Fig. S3), suggesting that collagen is produced but that proper CC organization depends on intact CA formation.

Together, these results indicate that Shroom3 is required for CA formation and contractile function in PCs, thereby maintaining normal lamellar architecture.

### Shroom3 regulates pillar cell arrangement and lamellar architecture in zebrafish gills

Having established that Shroom3 controls CA formation and promotes PC constriction, we next asked how these defects affect lamellar architecture at the tissue level. To address this, we performed whole-organ clearing of adult zebrafish gills using the *fgf10b:nEOS* reporter line, in which PCs express nuclear Eos fluorescent protein (Fig. 4A,B; Fig. S4A) ^25,30^. To minimize anatomical variability, we restricted our analysis to the dorsal region of the first gill arch (Movie 1).

**Fig. 4.**
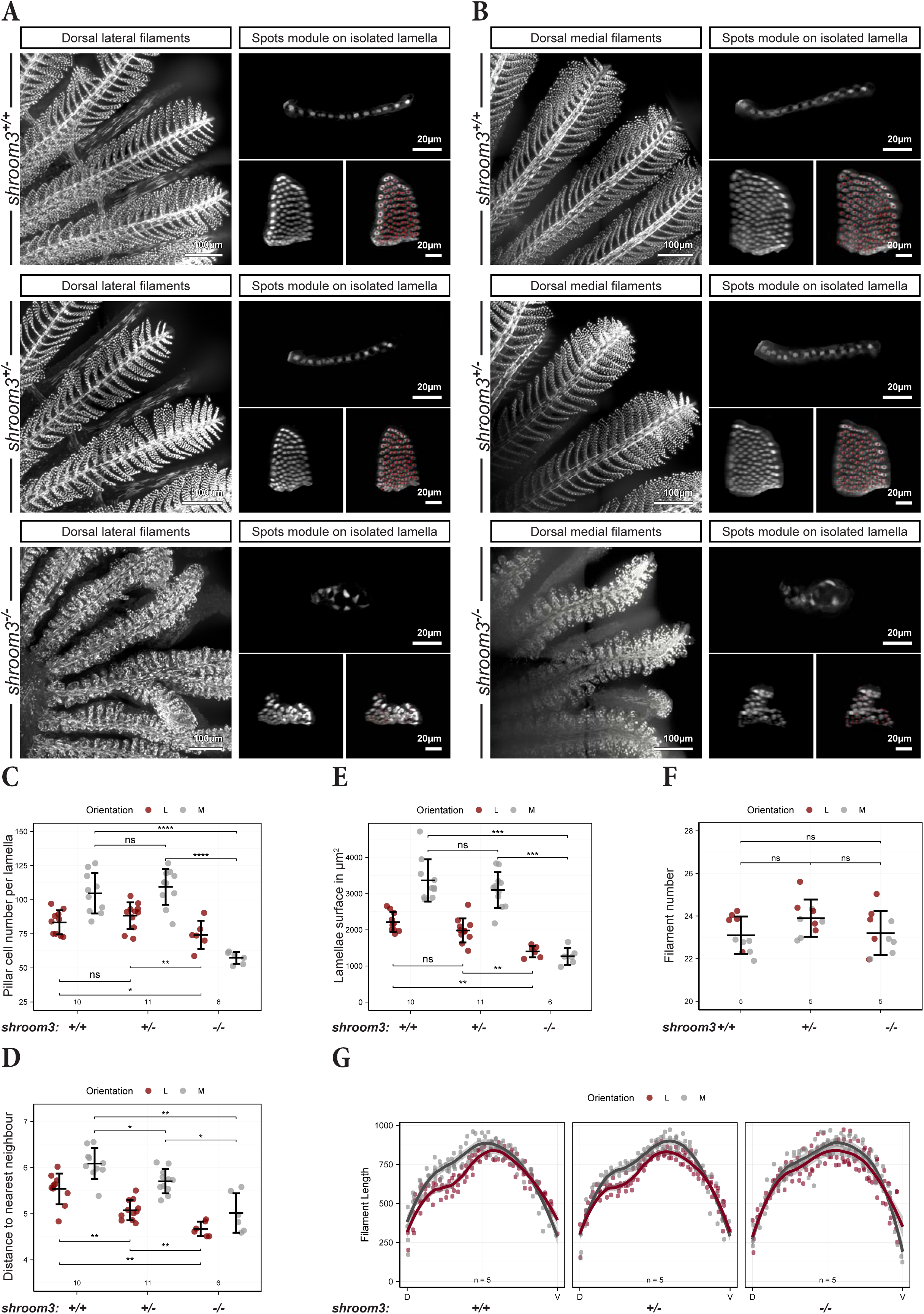
Shroom3 regulates pillar cell arrangement and lamellar architecture in zebrafish gills. (A-B) MIPs of cleared *fgf10b:nEOS* gills showing lateral (A) and medial (B) filaments in the dorsal region of a gill arch (left panel) across all *shroom3* genotypes as indicated. The right panels show representative single lamellae digitally isolated in Imaris with and without the Imaris Spots module (bottom), together with the corresponding orthogonal view (top). (C-E) Quantification of PC number (C) and PC spacing measured as the distance to the nearest neighbor (DTNN) (D), in lateral and medial filaments from the dorsal third of the gill arch across the indicated s*hroom3* genotypes. Each data point represents the mean value from three lamellae from one fish. The lamellar surface (E) was calculated using a mathematical model (see methods) based on PC number (C) and PC density (D). (F) Quantification of lateral and medial filament number from the dorsal region in the first gill arch across all *shroom3* genotypes. Each data point represents the total number of filaments in one hemibranch. (G) Length of lateral and medial filaments along the dorso-ventral axis of the first gill arch across all *shroom3* genotypes. Each data point represents one filament. (C-F) Error bars indicate mean ± SD. Two-tailed Mann-Whitney U-test: \**P* ≤ 0.05, \*\**P* ≤ 0.01, \*\*\**P* ≤ 0.001, ns = not significant. (C-G) Numbers below each panel indicate the number of animals analyzed (n) per genotype.

Consistent with previous work ^30^, lamellae from lateral filaments contained fewer PCs than those from medial filaments, and the cells were more densely packed (Fig. 4A-D). While the overall gill morphology and the total number of PCs per lamella were unchanged in *shroom3^+/-^* mutants, the distance to the nearest neighbour (DTNN) was significantly reduced in both filament types compared to wild-type siblings, indicating increased PC density. Notably, the previously reported medio-lateral difference in PC density was preserved in *shroom3^+/-^* mutants (Fig. 4D) ^30^. This increase in density, in the absence of a change in cell number, was accompanied by an overall reduction in lamellar size (Fig. 4E).

The severely malformed lamellae of *shroom3^-/-^* mutant gills showed a significant reduction in PC number per lamella in both lateral and medial filaments compared with wild-type and heterozygous siblings (Fig. 4C). Surprisingly, medial lamellae were more affected, reversing the medio-lateral asymmetry. PCs in *shroom3^-/-^* lamellae were densely packed and irregularly distributed (Fig. 4A,B,D; Fig. S3), lacking the regular spacing observed in control siblings (Fig. 4A,B; Fig. 1E; Fig. S3). As a result, lamellae lost their characteristic sheet-like structure and instead formed smaller, irregular outgrowths (Fig. 4A,B; Movie 2).

Despite this severe lamellar disorganization, overall filament architecture was largely preserved in *shroom3^-/-^* mutants, with similar filament number (Fig. 4F), distribution and length (Fig. 4G; Fig. S4B) compared with siblings. Occasional distal branching events were observed (Fig. S4A).

Collectively, these data show that Shroom3 is required not only for PC constriction but also for proper PC spacing, thereby preserving the architecture of the lamellar vasculature.

### Early lamellar defects in *shroom3* mutants are associated with chronic hypoxia

Having established that Shroom3 regulates CA formation and constriction in adult zebrafish, we next asked when these defects arise during development. In a recent study, we showed that lamellar formation begins at approximately 4-5 dpf ^30^. At 6 dpf, lamellae were actively forming and analysis of *fli1a:GFP* larvae revealed GFP-positive PCs within newly emerging lamellae (arrowhead in Fig. 5A). Concanavalin A staining marked CCs within these *fli1a:*GFP-positive PCs in both wild-type and *shroom3^+/-^*siblings (arrows in Fig. 5A), indicating that CC formation occurs concomitantly with early PC differentiation (Fig. S5). In contrast, no CCs were detectable in *shroom3^-/-^* larvae at this stage and lamellae already appeared swollen and malformed by 6 dpf (Fig. 5A, lower panel).

**Fig. 5.**
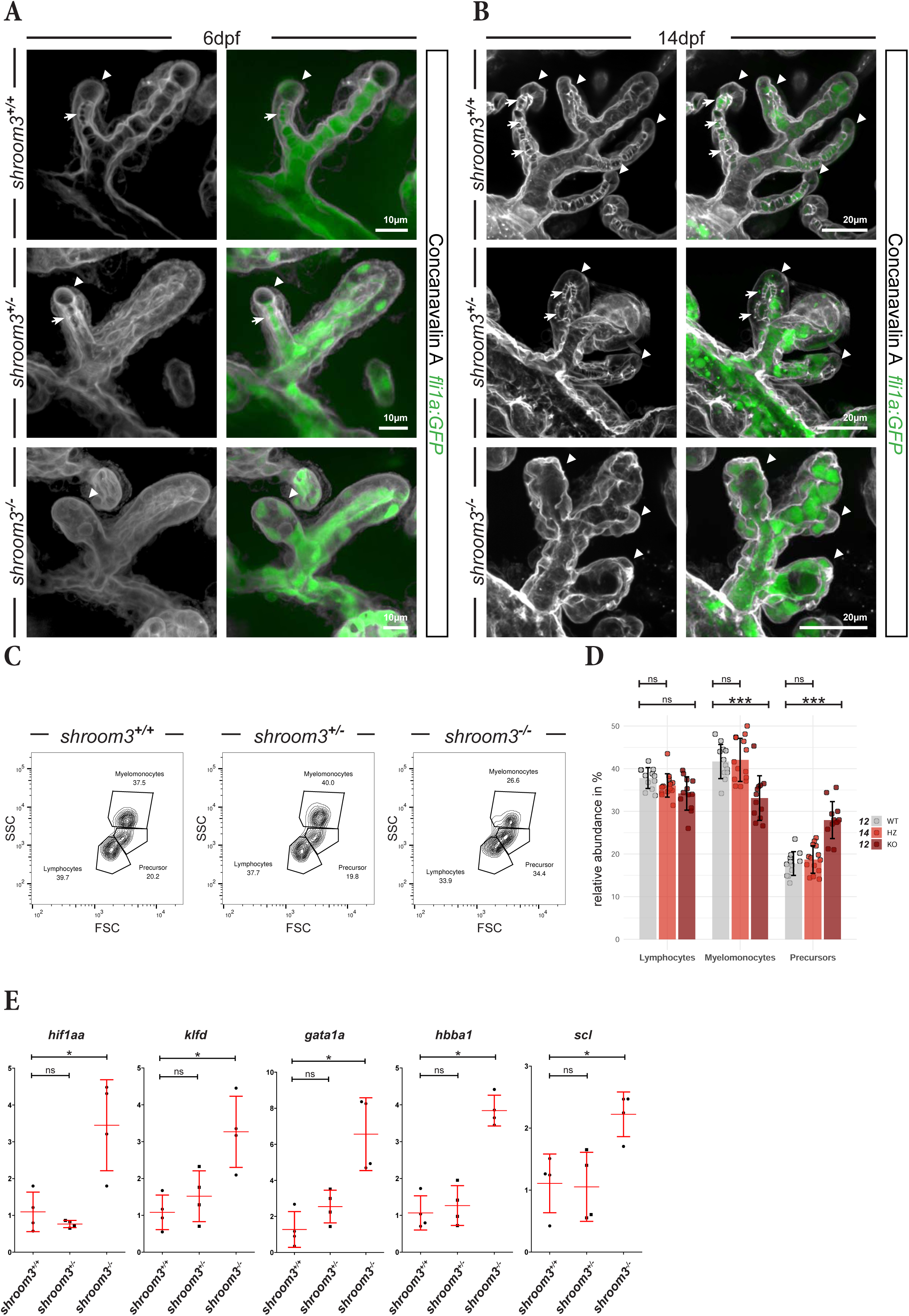
Early lamellar defects in *shroom3* mutants are associated with chronic hypoxia. (A-B) Representative MIPs of gill filaments from 6-dpf (A) and 14-dpf (B) *fli1a:GFP* larvae of the indicated *shroom3* genotypes labeled with Concanavalin A to visualize collagen columns. Arrowheads indicate lamellae; arrows indicate collagen columns between pillar cell bodies. (C) Flow cytometry analysis of whole kidney marrow (WKM) from the indicated *shroom3* genotypes. Representative scatter plots show gated lymphocyte, myelomonocyte and hematopoietic precursor populations. (D) Quantification of the relative proportions of WKM lymphocyte, myelomonocyte and hematopoietic precursor populations in the indicated *shroom3* genotypes (n≥12 fish per genotype). (E) Relative expression of erythropoiesis- and hypoxia-related genes in adult WKM across the indicated *shroom3* genotypes, quantified by RT-qPCR and normalized to wild type. Three mesonephroi were pooled per sample; four pooled samples (12 fish total) were analyzed per genotype. (D,E) Error bars indicate mean ± SD. Two-tailed Mann-Whitney U test: \**P* ≤ 0.05, \*\*\**P* ≤ 0.001, ns, not significant.

By 14 dpf, additional lamellae had formed and more PCs exhibited fully established CCs in both wild-type and heterozygous fish (arrows in Fig. 5B). The lamellae of *shroom3^-/-^* mutants had further increased in size, resulting in even greater architectural disruption and progressively displacement of PCs towards the periphery (Fig. 5B, lower panel), likely reflecting the combined effects of missing CCs and hemodynamic load.

This early onset of the phenotype prompted us to test whether the lamellar defects in *shroom3^-/-^*mutants compromise oxygen uptake and induce chronic hypoxia. Because erythropoiesis is upregulated under hypoxic conditions ^31^, we analyzed hematopoietic cell populations in whole kidney marrow (WKM) by flow cytometry using established light-scatter parameters ^32^ (Fig. 5C). *shroom3^-/-^*fish exhibited a significant reduction in myelomonocytes and a trend toward fewer lymphocytes (Fig. 5D). Consistent with our hypothesis, they also showed elevated levels of hematopoietic precursors, suggesting a compensatory response to reduced oxygen availability (Fig. 5D). RT-qPCR analysis of renal tissue further revealed significant upregulation of the erythropoiesis-related genes *gata1a*, *hbba, scl*, and *klfd*, as well as the hypoxia marker *hif1aa*, in homozygous mutants (Fig. 5E).

Together, these findings show that loss of *shroom3* disrupts lamellar architecture from the onset of lamellar development and is associated with a systemic erythropoietic response consistent with chronic hypoxia, thereby linking PC morphogenesis to oxygen homeostasis.

### Shroom3 deficiency increases phosphorylated myosin levels within the pillar cell contractile apparatus

Shroom3 promotes actomyosin contractility by physically interacting with and recruiting Rho-associated kinase (ROCK) to sites of apical constriction in epithelial cells, thereby promoting NMII activity through ROCK-dependent phosphorylation ^20,33,34^. We therefore tested whether Shroom3 regulates PC longitudinal constriction through a similar mechanism.

We stained gill cryosections with Phalloidin and an antibody against phosphorylated myosin light chain (pMLC) to visualize active NMII. Consistent with Fig. 2C, phosphorylated myosin localized to the Phalloidin-positive CAs surrounding PC nuclei, indicating enrichment of active myosin at the CA (Fig 6A,B; Fig. 2C). The staining again highlighted regional differences in PC constriction state and was consistent with the previously observed increased constriction in peripheral PCs compared to central PCs, as peripheral PCs exhibited elevated phospho-myosin fluorescence intensity at the CA (Fig. 6A,C; Fig. S2). Unexpectedly, quantification of mean fluorescence intensity at the CA revealed a significant increase in phosphorylated myosin levels in *shroom3^+/-^* mutants compared with wild-type siblings (Fig. 6A,B,D). This suggests that partial loss of Shroom3 triggers compensatory upregulation of myosin activity in response to reduced CA number and altered PC morphology, possibly to preserve lamellar integrity (Fig. 3B).

**Fig. 6.**
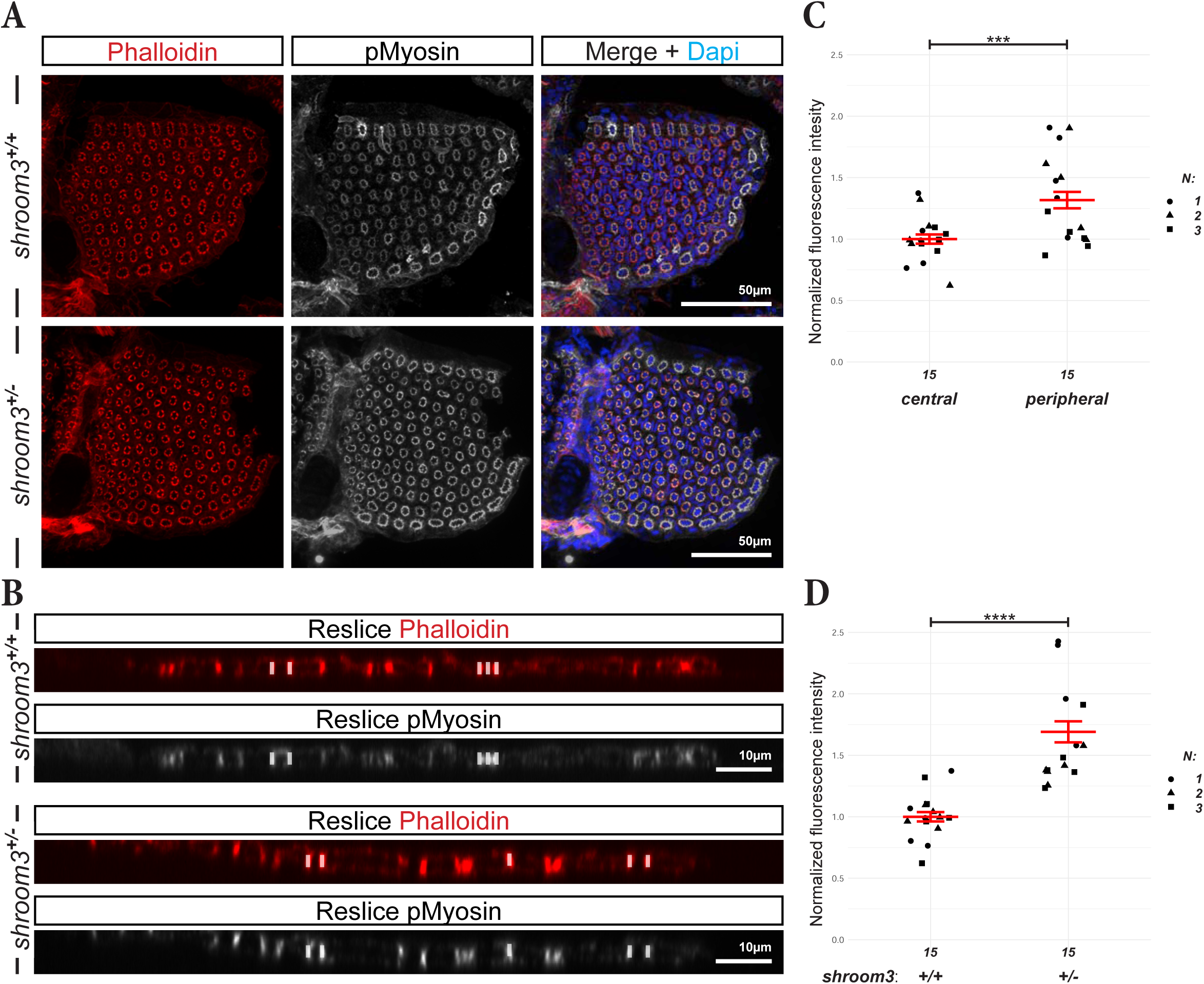
Shroom3 deficiency increases phosphorylated myosin levels within the pillar cell contractile apparatus. (A) MIPs of cryosections from adult gill arches labeled with phalloidin and DAPI and immunostained for phospho-myosin in shroom3^+/+^ and shroom3^+/-^ fish. (B) Orthogonal projections of the stacks in (A) showing cross-sections through lamellae. Genotypes and channels are indicated. White bars indicate representative regions of interest selected in the phalloidin channel and used for blinded quantification of fluorescence intensity at the CA in the pMyosin channel. (C, D) Quantification of p-Myosin mean fluorescence intensity at the CA of peripheral and central PCs in *shroom3^+/+^* fish (C) and in shroom3^+/+^ versus shroom3^+/-^ fish (D). Each data point represents the mean value from 90 measurements obtained from three different lamellae of one fish (30 measurements per lamella). Values were normalized to the wildtype mean. The experiment was performed independently three times with five fish per genotype (N=3). Error bars represent mean ± SD. Statistical significance was assessed using a two-tailed Mann-Whitney U-test: ***P ≤ 0.001; ****P ≤ 0.0001.

### Vinculin b regulates contractile apparatus formation and pillar cell constriction

Given the central role of Shroom3 in CA organization and function, we next investigated mechanosensitive proteins that could transmit CA-generated forces and thereby contribute to PC function. Vinculin (Vcl) is a mechanosensitive adaptor that localizes to adhesion complexes and undergoes force-dependent conformational changes that promote actin recruitment ^35^. Because Vinculin localized to the CA in PCs (Fig. 2E), it represented a strong candidate for mediating mechanical force transmission in this system. Supporting this, a recent study using scRNA-seq analysis revealed transcriptional similarities between zebrafish PCs and mammalian aerocytes ^36^, including specific enrichment of Vinculin expression. Among the two zebrafish Vinculin paralogs *vclb* was identified as the predominantly expressed form in PCs, motivating subsequent functional analysis of *vclb* ^27^.

In contrast to the severe swelling and broad disorganization observed in *shroom3^-/-^* mutants, cleared *vclb^-/-^* gills exhibited a milder phenotype characterized by localized swelling at the base of lamellae (arrowheads in Fig. 7A), near the blood flow entry and exit sites from the afferent and efferent filament arteries.

**Fig. 7.**
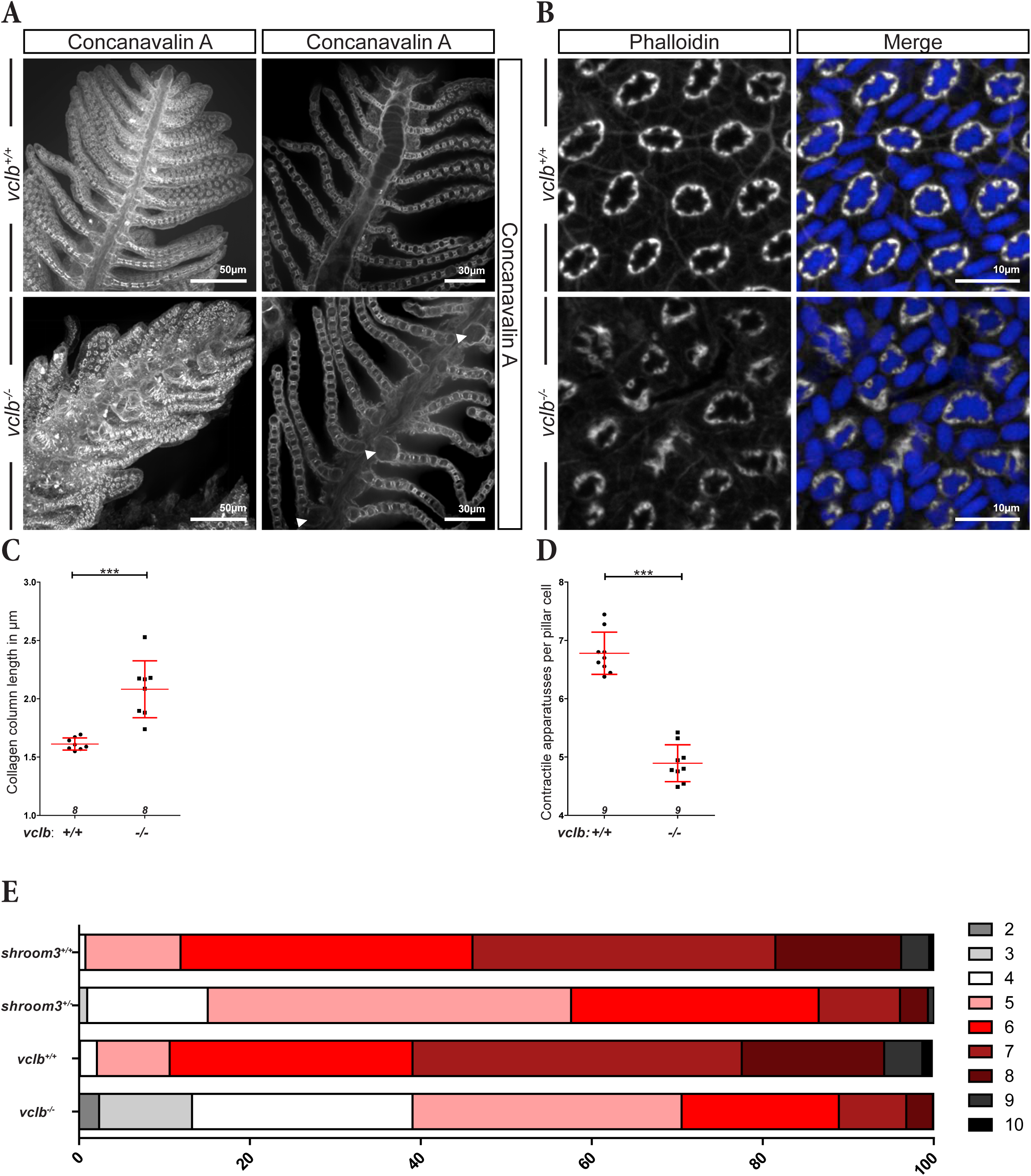
Vinculin b regulates contractile apparatus formation and pillar cell constriction. (A) MIPs of ECi-cleared gills in *vclb^+/+^* and *vclb^-/-^* fish stained with Concanavalin A to label collagen columns (left). MIPs of smaller Stacks (2,5μm) are shown on the right, highlighting locally enlarged VS (arrowheads). (B) MIPs of cryosections from adult gill arches labeled with phalloidin and DAPI in *vclb^+/+^* and *vclb^-/-^* fish. (C, D) Quantification of CC length (C) and CA number per PC (D) in *vclb^+/+^* and *vclb^-/-^* fish. Error bars indicate mean ± SD. Statistical significance was assessed using a two-tailed Mann-Whitney U: \*\*\**P* ≤ 0.001. Each data point represents the mean value of 90 PCs from one fish. Numbers at the bottom of each panel indicate the number of fish analyzed (n). (E) Percentage bar graph showing the relative distribution of CA number per PC in *shroom3^+/+^*, *shroom3^+/-^, vclb^+/+^* and *vclb^-/-^* fish. Values are derived from the quantifications shown in Fig. 3B and Fig. 7D.

This spatially restricted phenotype suggested that PCs locally collapse in regions exposed to elevated hemodynamic load, consistent with a role for Vinculin in force transmission. To test whether PC constriction was impaired, we measured CC length in intact PCs in *vclb^-/-^* mutants (Fig. 7A,C). CCs were significantly longer in *vclb^-/-^* mutants than in wild-type siblings, indicating reduced PC constriction. Notably, the degree of CC elongation was comparable to that observed in *shroom3^+/-^* mutants (Fig. 3F). However, although *shroom3^+/−^* mutants displayed elongated CCs, they did not exhibit localized lamellar swelling, suggesting that CC elongation alone is insufficient to explain this phenotype. We therefore asked whether additional defects in CA organization might account for the localized lamellar swelling seen specifically in *vclb^-/-^* mutants. Quantification of CA number revealed a significant reduction in *vclb^-/-^* PCs, similar to *shroom3^+/-^* mutants (Fig. 7B,D; Fig. 3B). Notably, *vclb^-/-^* PCs contained, on average, 0.5 fewer CAs per cell than *shroom3^+/-^*(Fig. 7D). This difference was even more pronounced at the population level: while only ∼15% of PCs in *shroom3^+/-^*mutants contained fewer than five CAs, this fraction increased to ∼40% in *vclb^-/-^* mutants, with some cells forming as few as two CAs (Fig. 7E). In addition, the CAs in *vclb^-/-^*PCs appeared malformed and unevenly distributed, indicating compromised structural organization (Fig. 7B).

Together, these findings identify Vinculin b as a key regulator of CA formation and PC constriction. Loss of *vclb* reduces CA number and impairs constriction, resulting in partial disruption of lamellar architecture and an intermediate phenotype between *shroom3^+/-^* and *shroom3^-/-^*mutants. These findings suggest that Shroom3-dependent actomyosin contractility and Vinculin-mediated force transmission act in concert to stabilize CA architecture in PCs.

## Discussion

### Structural organization and heterogeneity of pillar cells

PCs are the core structural and mechanical elements of the gill lamellae, where they maintain vascular architecture and tissue integrity. Yet their functional organization has remained incompletely understood. Previous studies have described aspects of the PC architecture in pufferfish (*Takifugu rubripes)* ^21,23^, examined their developmental origin by lineage tracing of PC progenitors ^25,26^ and characterized their transcriptional profile in zebrafish ^27^. However, a systematic structural and quantitative analysis of PC architecture and function in zebrafish has been lacking, with only partial ultrastructural descriptions available from electron microscopy studies ^24,27,30^.

Although the overall organization of zebrafish PCs resembles that reported in pufferfish ^21,23^, our observations provide a more detailed view of PC architecture and reveal additional features not described previously. In particular, we identified a continuous ZO-1-positive tight junctions (TJ) network that seals not only each CA, but also the VS formed between adjacent PCs. Its colocalization with the outer actin ring suggests that this junctional network may contribute not only to barrier function but also to mechanical coupling between neighboring PCs, thereby stabilizing the vascular compartment. Furthermore, we provide evidence consistent with integrin-mediated adhesion linking the intracellular actomyosin-rich CA to the extracellular CCs. The localization of Itgb1b together with Vinculin and Paxillin is reminiscent of focal adhesion complexes that mediate physical coupling between cells and the ECM ^5,37^. We therefore propose that PCs are anchored to the CCs via a focal adhesion–like machinery that enables contractile forces generated by the CA to be transmitted to the extracellular scaffold.

Beyond this structural characterization, our quantitative analyses uncover an unexpected degree of heterogeneity within the PC population. By systematically quantifying CA number and assessing constriction state via CC length, we identified differences between peripheral and central PCs within individual lamellae. Peripheral PCs contained more CAs and shorter CCs, consistent with a higher degree of longitudinal constriction, whereas central PCs displayed fewer CAs and longer CCs. This spatial pattern likely reflects differential mechanical demands across the lamella and the need for increased structural stability near the OMC, which is exposed to higher hemodynamic stress ^27,38,39^. We also quantified PC number and spatial arrangement, as well as lamellar surface area and filament number and length. In wild-type and heterozygous fish, these values, including the medio-lateral asymmetry were consistent with our previous analysis ^30^. In contrast, *shroom3^-/-^* lamellae showed a strong reduction in PC number, especially in medial filaments, raising the possibility that severe lamellar disruption feeds back on the homeostatic stem cell niche that supplies new PCs, at least in medaka ^40^.

Together, these findings establish PCs as highly specialized structural and contractile units of the lamella and provide a quantitative framework for their organization in zebrafish. This detailed characterization should facilitate future studies of PC architecture, their interaction with the ECM via CCs, as well as their spatial arrangement and mechanical function. This may be particularly relevant in light of recent evidence highlighting similarities between PCs and mammalian lung aerocytes ^27^.

### Shroom3 promotes pillar cell constriction by regulating contractile apparatus formation

Shroom3 is best known for its role in apical constriction in epithelial tissues ^7,12–14,18,20,33,41^. Our findings extend this function to gill PCs, a cranial neural crest-derived lineage, thereby broadening the developmental and cellular contexts in which Shroom3 regulates cell shape. This is consistent with the role of SHROOM3 in morphogenetic force generation in cardiac neural crest cells during mouse development and congenital heart disease ^42,43^.

The capacity of PCs to constrict was proposed decades ago based on the presence of actomyosin-rich contractile structures within these cells ^22,44^. Additional support came from physiological studies in Atlantic cod, another actinopterygian species, showing that injection of the vasoconstrictive hormone endothelin type 1 increases PC diameter ^29,39,45^. These findings indicate that CA constriction alters overall PC morphology making them wider and shorter, and also modifies VS morphology. However, these studies were largely descriptive, and a quantitative understanding of the underlying cellular and molecular mechanisms has remained lacking. Our data now provide such a quantitative framework by linking CA organization to PC constriction and by identifying Shroom3 and Vinculin b as key regulators of this process.

Several lines of evidence support a model in which CA number is a major determinant of longitudinal PC constriction. Across individual PCs, higher CA number correlated with a more constricted state, reflected by a larger cell body area in cross-section and shorter CCs, whereas cells with fewer CAs exhibited smaller body areas and elongated CCs, consistent with reduced longitudinal constriction. This relationship was also reflected spatially, as peripheral PCs contained more CAs and were more strongly constricted than central PCs. Notably, peripheral PCs also exhibited increased phospho-myosin fluorescence intensity, raising the possibility that CA number and/or constriction state is associated with NMII activation. Together, these observations suggest that CA number contributes directly to the extent of PC constriction and thereby to cell shape. They further raise the possibility that CA organization is regionally adapted to local mechanical demands across the lamella.

Analysis of *shroom3^+/-^* mutants further supports this interpretation. Compared to wild-type siblings, PCs in *shroom3^+/-^* fish exhibited fewer CAs and longer CCs, indicating impaired constriction and supporting the idea that Shroom3 promotes CA assembly. At the mechanistic level, Shroom3 may contribute to this process through its known ability to associate with, and potentially recruit, actin via its ASD1 domain ^7,34,46^. In PCs, this could promote the assembly, stabilization, or spatial positioning of individual CAs. Whether Shroom3 primarily acts by recruiting pre-existing actin filament or by stabilizing newly assembled actomyosin structures remains to be determined. Based on ultrastructural observation, CCs were proposed to form during PC differentiation through the enclosure of extracellular collagen ^38^. Our findings support this hypothesis (Fig. S5) and raise the possibility that the enclosure depends on proper formation of the CA, including the autocellular junctions. In this context, the impaired assembly of CAs observed in *shroom3* mutants may compromise CC formation, ultimately leading to lamellar instability and the accumulation of CC remnants observed in *shroom3^-/-^*fish (Fig. S3).

### Shroom3 couples contractile apparatus organization to effective pillar cell constriction

Our data place Shroom3 upstream of CA architecture and support the view that proper CA formation is required for effective PC constriction. Yet, *shroom3^+/-^*PCs remained less constricted than wild-type PCs when comparing cells with the same number of CAs (Fig. 3C’). This indicates that Shroom3 not only promotes CA formation but also contributes to their contractile output and efficiency.

Given the well-established role of Shroom3 in promoting NMII activation through recruitment of ROCK1/2 ^20,34^, we initially expected reduced NMII phosphorylation in *shroom3^+/-^* PCs as an additional mechanism underlying the constriction defect. Unexpectedly, however, *shroom3^+/-^* PCs displayed increased phosphorylated NMII. One possible explanation is that this reflects a compensatory response, whereby PCs attempt to counteract reduced CA number by enhancing NMII activation at the remaining contractile structures. In support of this idea, compensatory kinase activity has been described in other systems. For example, in human detrusor smooth muscle, age-associated downregulation of ROCK2 is accompanied by increased expression of myosin light chain kinase (MLCK) ^47^, one of the major kinases that phosphorylate the myosin regulatory light chain ^48^. In addition, MLCK and ROCK differ in their spatial contributions to myosin activation, with MLCK preferentially targeting peripheral stress fibers and ROCK more strongly regulating centrally located actomyosin structures ^19,49^. By analogy, partial loss of *shroom3* in heterozygous mutants could reduce local ROCK recruitment without abolishing myosin phosphorylation in PCs, because other kinases such as MLCK may compensate. In this scenario, the elevated phospho-myosin signal in *shroom3^+/-^* PCs would reflect a compensatory activation state that remains insufficient to restore wild-type constriction, as indicated by the reduced PC cell body area and increased CC length in *shroom3^+/-^*mutants.

Another possibility is that Shroom3 contributes to PC constriction by supporting structural coupling between the actomyosin-rich CA and the extracellular CCs. Shroom3 can physically interact with actin and actin-binding proteins ^7,33,34^, raising the possibility that partial loss of Shroom3 compromises the mechanical integrity or force transmission capacity of the CA. In such a scenario, contractile forces may still be generated but transmitted less efficiently to the extracellular matrix. This interpretation is consistent with the established role of SHROOM3 in maintaining the glomerular filtration barrier ^15,17,18^, and with recent evidence that *SHROOM3* deficiency in podocytes results in the downregulation of focal adhesion-associated components such as Paxillin, Talin1 and integrins ^50^. Together, these findings suggest that Shroom3 is required not only for proper CA formation but also for the mechanical coupling allowing the conversion of contractility into effective cell constriction.

### Vinculin b may reinforce force transmission and stability at the contractile apparatus-matrix interface

The localization of Vinculin b at the CA, together with its requirement for maintaining lamellar integrity, particularly in regions likely exposed to elevated hemodynamic stress, is consistent with the well-established role of Vinculin as a mechanosensitive component of adhesion complexes ^51–55^. Based on its spatial association with the actomyosin-rich CA and with Itgb1b-positive adhesion sites, we propose that Vinculin b contributes to the mechanical coupling between the intracellular contractile machinery and the extracellular CCs (Fig. S1C-E, schematic). In this model, increased blood flow-associated forces could promote Vinculin b activation and conformational opening, thereby enabling recruitment of additional F-actin and strengthening the CA-CC linkage ^54,56,57^. Such force-dependent reinforcement would help sustain CA organization, stabilize lamellar architecture under load, and improve the efficiency with which contractile forces are transmitted to the extracellular scaffold. We propose that Shroom3 and Vinculin b play complementary roles in PC mechanics: Shroom3 promotes CA formation and contractile competence, whereas Vinculin b likely reinforces force transmission at the CA–matrix interface, thus contributing to the conversion of contractile activity into tissue-scale stability.

### A mechanical threshold model of the contractile apparatus underlies lamellar stability

Our data support a threshold model in which lamellar stability depends on a minimum number of functional CAs per PC. In this framework, each CA acts as a force-generating and load-bearing unit, and lamellar integrity is maintained only when enough CA/CCs are present to counteract hemodynamic pressure within the vascular space.

The graded phenotypes observed across genotypes are consistent with this model. In *shroom3^+/-^* fish, PCs contained fewer CAs and were less constricted than in wild-type siblings, yet lamellae remained largely intact. This suggests that CA number is not reduced below the level required to preserve overall tissue integrity, potentially aided by compensatory changes such as increased PC density and elevated myosin phosphorylation. Because this reduction affects PCs broadly across the lamella, it likely weakens the tissue at a global level. Thus, even increased myosin phosphorylation may be insufficient to restore effective constriction, because the same hemodynamic load must still be borne by a uniformly weakened PC network despite otherwise largely preserved tissue architecture.

In *vclb^-/-^* mutants, CA number and organization were more severely affected, and lamellae exhibited localized collapse, particularly in regions likely exposed to higher hemodynamic load. Finally, in *shroom3^-/-^* lamellae, CA formation was profoundly disrupted, CCs were absent, and the lamellar structure collapsed globally. Together, these phenotypes represent a continuum that correlates with the extent of CA loss and disorganization, suggesting that a minimum number of CA/CCs per PC is required to mechanically stabilize the lamellar architecture and counteract hemodynamic pressure. This model also provides a plausible explanation for the spatial pattern of collapse. In *vclb*^-/-^ gills, instability was most apparent near the afferent and efferent filament arteries, where local pressure is expected to be highest. By contrast, in *shroom3*^-/-^ mutants, the mechanical threshold appears to be breached throughout the lamella from the onset of development, preventing proper establishment of the supporting PC architecture and ultimately resulting in vascular expansion, lamellar ballooning, and chronic hypoxia.

Interestingly, this principle resembles a recent model of *Drosophila* wing morphogenesis, in which regularly spaced epithelial pillars counteract internal pressure during wing deployment and maintain a luminal space between neighboring cells ^58^. Together, our findings identify CA organization as a key determinant of lamellar mechanical integrity and support a model in which Shroom3- and Vinculin b-dependent CA architecture sets the mechanical threshold required for lamellar stability, thereby linking subcellular contractility to tissue-scale integrity.

## Methods

### Zebrafish lines and maintenance

All animal procedures were conducted in accordance with the guidelines of Goethe University Frankfurt and the German Animal Welfare Act and were approved by the German authorities (veterinary department of the Regional Board of Darmstadt). Zebrafish (*Danio rerio*) were raised and maintained under standard conditions ^59^. The following transgenic lines have been described previously: *Tg(kdrl:Hras.HRAS-mCherry)^s896^*(*kdrl:mCherry*) ^60^, *Tg2*(*NLS-Eos*)*^el865^* (*fgf10b:nEOS*) ^25^, *Tg*(*fli1a:EGFP)^y^*^1^ (*fli1a:GFP*) ^61^, *Tg(actb2:myl12.1-eGFP)^e2212^* (*myl12-GFP*) ^62^ have been described previously. To assess the contribution of Vinculin b, we used the *vclb^bns2^*^47^ mutant line, which carries a 16 bp deletion ^63^.

### Embryo Genotyping

Embryos, larvae or fin clips were lysed in 50 mM NaOH at 95°C for 1 h. Samples were neutralized by adding 1:10 of the reaction volume of 1 M Tris-Cl (pH 8.2). Following neutralization, 2 μl of the genomic DNA lysate was used as template for PCR-based genotyping. Primers for genotyping are presented in Table S1. Genotyping PCR products were resolved on 4% agarose gels.

### In situ hybridization

Whole-mount colorimetric *in situ* hybridization (ISH) was performed according to standard procedures^64^. To improve the signal-to-noise ratio for imaging, dissected gill arches were dehydrated in 100% ethanol for 1 h, cleared in ethyl cinnamate (ECi), and imaged.

### Hybridization Chain Reaction (HCR)

Dissected gills were fixed in 4% paraformaldehyde (PFA) in PBS overnight at 4°C and processed as for conventional ISH up to the proteinase K digestion and postfixation steps. HCR was subsequently performed according to the manufacturer’s instructions (Molecular Instruments). HCR probe were designed and synthesized by Molecular Instruments based on the submitted target gene. Because erythrocytes within the lamellae exhibited strong autofluorescence, background fluorescence was computationally subtracted using Fiji.

### Cryosections and gill dissection

Adult fish were euthanized by hypothermic shock in 4°C system water according to the protocol approved by the local authorities. Gills were then dissected directly in 4% PFA in PBS and fixed for 2 h at room temperature (RT) or overnight at 4°C. Samples were washed three times for 5 min in PBS and cryoprotected at 4°C overnight in 30% sucrose prepared in ddH20 until they sank. Gills were then oriented as desired in a 1.5 mL reaction tube containing a 1:1 mixture of 30% sucrose and OCT embedding medium (CellPath), flash-frozen in liquid nitrogen and sectioned at 18 µm using a CM3050 S cryostat (Leica). Cryosections were collected on SuperFrost Plus adhesion slides (Epredia) and stored at −80°C until further use.

### Immunofluorescence on cryosections

Cryosections were washed three times for 5 min in PBS and encircled with a hydrophobic barrier pen (Science Services). After blocking for 2 h at RT in 10% normal goat serum (NGS), sections were incubated overnight at 4°C with one of the following primary antibodies diluted at 1:200 in 2% NGS: mouse anti-Paxillin (BD Biosciences, 610051), mouse anti-Vinculin (Sigma, SAB4200729), rabbit anti-phospho-MLC2 (Cell Signaling Technology, 3671), mouse anti-ZO1 (Thermo Fisher Scientific, 33-9100). After washing 3 times in PBS for 5 min, sections were incubated 2 h at RT with the appropriate cross-adsorbed secondary antibodies diluted at 1:400 in 2% NGS: goat anti-rabbit Alexa Fluor 647 (Thermo Fisher Scientific, A21244), goat anti-mouse Alexa Fluor 647 (Thermo Fisher Scientific, A21235). Where relevant indicated, Phalloidin Alexa Fluor 594 (2 U/mL), DAPI (1 µg/mL) and Concanavalin A-Alexa Fluor 647 (5 µg/mL) (all Thermo Fisher Scientific) were added for 30min. Sections were mounted in Mowiol mounting medium.

### Ethyl cinnamate-based clearing procedure

Gills were dissected in 4% PFA in PBS under the hood and fixed for 2 h at RT. Samples were washed three times for 30 min at 4°C in ice-cold PBS containing 0.3% Triton X-100. Where indicated, gills were incubated with either Concanavalin A-Alexa Fluor 647 (4 U/mL; Thermo Fisher Scientific) or rabbit anti-EosFP antibody (1:300; Novus Biologicals, NBP3-05557) diluted in 2% NGS for approximatively 60 h at 4°C. Samples were subsequently washed overnight in PBS.

Samples were dehydrated in a graded ethanol series (50%, 70% and 2× 100% ethanol, 4h per step) using a TP1020 autostainer (Leica). Samples were then incubated in ECi (Sigma-Aldrich, 112372) for 2 h prior to imaging. For imaging, samples were transferred onto a slide, immersed in ECi, and gently sealed with a coverslip.

### Spinning-disc microscopy

Fluorescence imaging was performed using a fully motorized Nikon Ti spinning-disc confocal microscope controlled with NIS-Elements 5.0 software.

### Electron microscopy

Dissected gills were fixed in 2.5% glutaraldehyde and 4% PFA in 0.1 M cacodylate buffer (pH 7.2) for 2 h at RT. For TEM, samples were washed twice in 0.1 M cacodylate buffer containing 2% sucrose, post-fixed in 1% reduced osmium tetroxide, dehydrated and embedded in Araldite resin. Ultrathin sections (50 nm) were cut using an ultramicrotome (Leica), and ribbons of sections were collected on Formvar-coated copper slot grids. Sections were contrasted with 5% uranyl acetate in methanol/water followed by lead citrate. Micrographs were acquired using a TEM 900 transmission electron microscope (Zeiss) operated at 80 kV in bright-field mode and equipped with a Tröndle 2K camera.

For SEM, samples were critical-point dried after fixation, followed by gold sputter coating using the BALTEC MED 020 coating system. Imaging was performed using a Hitachi S-4500 SEM operated at 10-15 kV.

### Flow Cytometry analysis of whole kidney marrow

Adult fish were euthanized by hypothermic shock in 4°C system water. Following a ventral incision, the mesonephric kidney was dissected using forceps and mechanically dissociated by gentle pipetting approximatively 25 times in 600 µl FACS-Buffer (2% fetal calf serum (FCS) in PBS). The resulting cell suspension was passed twice through a 40 µm cell strainer (pluriSelect). Cells were stained with DAPI (1:1000 dilution in a final volume of 1 mL) for 10 min to exclude dead cells and debris ^32^. Following centrifugation (1500 rpm for 5 min), cells were resuspended in 550 µl FACS-Buffer. 50.000 events per sample were collected using a BD LSRFortessa Cell Analyzer. Total cells were gated based on forward scatter (FSC) and side scatter (SSC), and erythrocytes and debris were subsequently excluded (Fig. S6). Within this gate, myelomonocytes, precursor cells, and lymphocytes were identified by their distinct FSC and SSC profiles using contour plots. Relative number was calculated as the percentage relative to total events after exclusion of erythrocytes and debris.

### RNA extraction, cDNA synthesis and RT-qPCR

Total RNA was isolated using TRIzol (Invitrogen) according to the manufacturer’s instructions. One complete gill or three pooled mesonephroi were homogenized in 500 µl TRIzol using a pestle followed by passage through a blunt-end syringe needle. cDNA was synthesized using the iScript cDNA Synthesis Kit (Bio-Rad). For each reaction, 500ng of total RNA were reverse-transcribed into cDNA and was diluted 1:5. 2µl of diluted cDNA was used per well.

Primers were designed using Primer-BLAST and Primer efficiency was determined as described previously ^65^. For primer efficiency assays, cDNA was serially diluted 1:2 to obtain a total of six concentrations (25, 12.5, 6.25, 3.125, 1.5625 and 0.78125 ng/µL), each analyzed in technical triplicates. Real-time PCR was performed using iTaq Universal SYBR Green Supermix (Biorad, 1725270) in a CFX Connect Real-Time PCR detection System (Biorad). Relative gene expression was calculated using the ΔΔCt method ^66^ with *rpl13a* as the reference gene. Primers used for RT-qPCR are listed in table S1.

### Quantification of filaments, pillar cells and lamellar surface area

Imaris software (Oxford Instruments, v.10.1.1) was used for three-dimensional image visualization, image processing, quantification and video rendering. Gill arches, filaments and lamellae were digitally segmented manually using the Imaris Surface module (Fig. 4A-B; Fig.S4A; Fig.S5). Filament number and height were quantified as previously described (Fig. 4F-G; Fig.S4) ^30^. PC number and DTNN were quantified using the *fgf10b:nEOS* transgene signal and the Imaris Spots module as previously described (Fig.4C-D) ^30^, except that a spot diameter of 3,5 µm was applied for *shroom3^+/+^* and *shroom3^+/-^*, whereas a diameter of 2 µm was applied for *shroom3^-/-^*gills. Lamellar surface area was calculated from PC number and DTNN measurements as previously described ^30^.

### Fluorescence intensity measurement

Phosphorylated myosin fluorescence intensity at the CA was quantified using a custom Fiji macro. For each image, reslices were generated prior to analysis. The macro was designed to enable blinded and reproducible measurements from defined linear regions of interest (ROIs) in microscopy images. ROIs were drawn as straight lines (width = 5 px) along the CA in the phalloidin channel, which served as the blinding channel (Fig. 6B). For each specimen, 30 measurements were performed on three different lamellae (90 measurements in total). The ROIs were then automatically transferred to the phosphorylated myosin channel for quantification. For each ROI, the macro generated a plot profile and recorded the corresponding mean fluorescence intensity.

### Contractile apparatus quantification and pillar cell body area measurement

CA number and PC body area were manually quantified on phalloidin-labelled gill cryosections. First, CAs were manually counted in a total of 90 PCs per specimen, consisting of 30 PCs from each of three images (Fig. 3B; Fig. S2B,C; Fig. 7C). PC body area was subsequently measured in the same cells used for the CA number (Fig. 3C). PCs were manually outlined using the polygon selection tool, and cell area was measured. All measurements were performed in Fiji.

### Quantification of collagen column length

CC length was quantified on whole Eci-cleared gill arches stained with Concanavalin A. For each specimen, three images of lamellar cross-sectional views were generated. Given the high variability of lamellar size along the dorso-ventral axis ^30^, analyses were restricted to the dorsal portion of the first gill arch to ensure data comparability. The first ten lamellae, starting from the distal tip of each filament, were considered to be part of the growth domain ^40^ and were therefore excluded from the analysis. In total, 90 PCs were analyzed per specimen, consisting of 30 PCs from each of three images. Three CCs were measured per PC using the Fiji line selection tool. The mean CC length was first calculated for each PC and subsequently averaged per specimen.

### Statistics and data visualization

Unless otherwise stated, GraphPad Prism (v.5) was used for data visualization, with preliminary data processing performed in Microsoft Excel. Statistical analyses were performed using two-tailed unpaired Mann-Whitney U tests). P values are reported in the corresponding figure panels (i.e.: \*\*\**P* ≤ 0.001).

Analyses presented in Figures 3C’; 4C-G; 5D; 6C-D and S4B were performed in R (v.4.3.1) using R studio (2025.9.2.418). Data wrangling was carried out using dplyr (v.1.1.3) and tidyverse (v.2.0.0). Data visualization was performed with ggplot2 (v.4.0.1) and statistical analyses were conducted using ggpubr (v.0.6.0) and rstatix (v.0.7.2). Unless otherwise stated, statistical comparisons were performed using the Wilcoxon rank-sum test (Mann-Whitney U test). Comparisons between two groups, the *stat_compare_means* function was used (ggpubr v.0.6.0), whereas pairwise comparisons across multiple groups were performed using pairwise Wilcoxon rank-sum tests with Benjamini-Yekutieli (BY) correction for multiple testing (rstatix v.0.7.2.). These non-parametric tests were chosen for their robustness to non-normal distributions, assuming similar distribution shapes across groups. P-values are reported in the figure panels (i.e.: \*\*\**P* ≤ 0.001).

## Acknowledgements

We thank Prof. Julien Bertrand for sharing his expertise on zebrafish kidney marrow hematopoiesis and Dr. Julien Rességuier for valuable insights into gill biology and methodology. We are also grateful to Prof. Ivica Grgic for providing access to the tissue-clearing setup. We thank our bachelor students, L. Schwahn and M. Beranek, for their help. Special thanks go to the animal care team for the excellent maintenance of the fish facility, and to M. Kamprad and M. Heyde for their valuable technical assistance. We also thank the GRADE Center SCALE-it, the Wilhelm Foundation, and the European Zebrafish Society for providing travel funding.

## Funding

This work was supported by the Clusterproject EnABLE, which is funded by the Hessisches Ministerium für Wissenschaft und Kunst (V.L.).

## Declaration of Interests

The authors declare no competing interests.

## Data and resource availability

All relevant data can be found within the article and its supplementary information.

## Declaration of generative AI and AI-assisted technologies in the manuscript preparation process

During the preparation of this work, the authors used AI-based language tools (ChatGPT, OpenAI, Claude) to assist in R code development and manuscript reformulation. The authors reviewed and edited the output as needed and take full responsibility for the content of the published article.

## Supplementary Figure legends

**Fig. S1.**
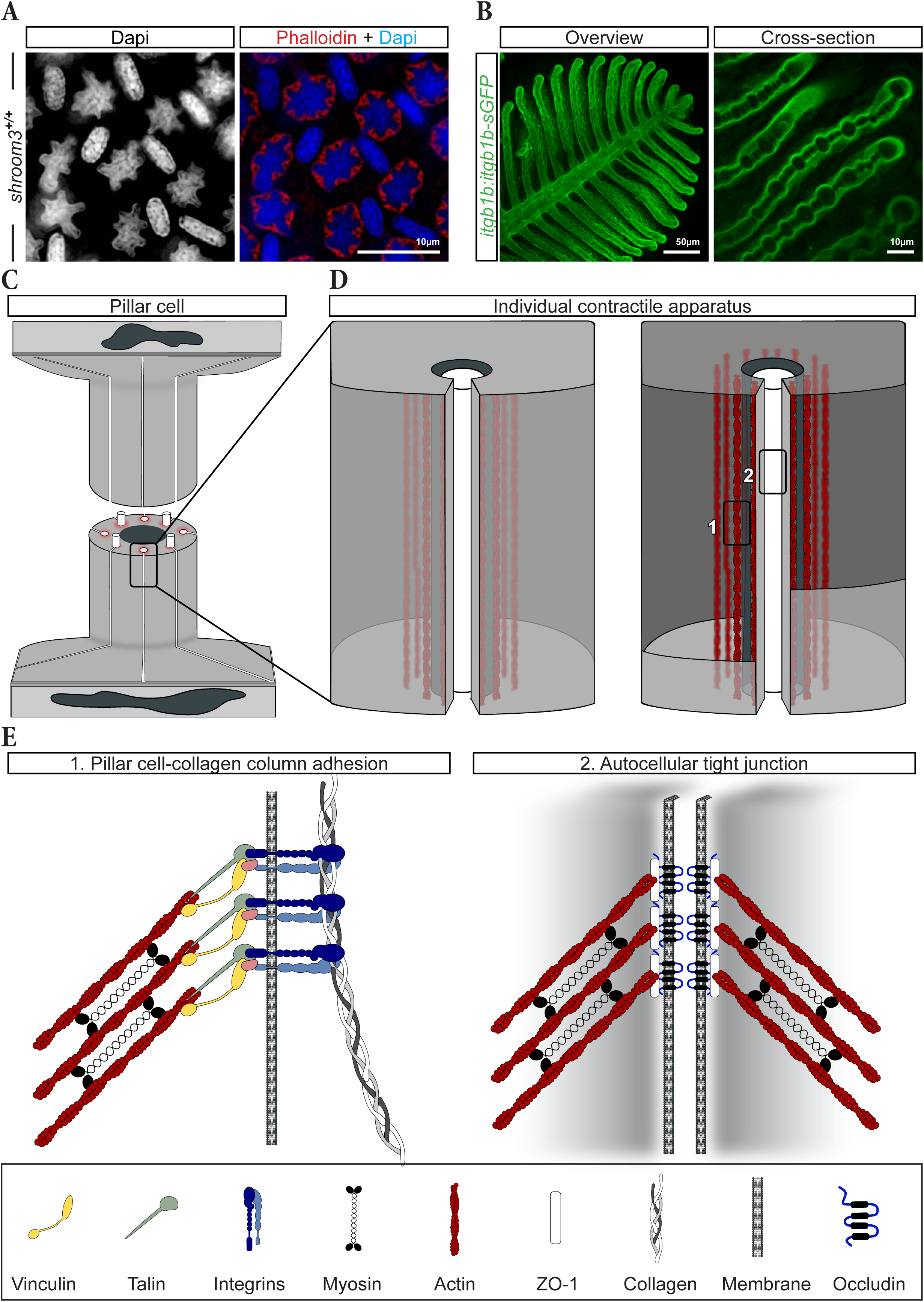
Itgb1b connects the collagen column to the contractile apparatus in gill pillar cells. (A) Cryosections stained with phalloidin to visualize the CA and with DAPI to label nuclei and imaged using Machine Intelligent Structured Illumination Microscopy (MI-SIM). (B) Representative MIP of a filament from dissected adult *Tg(itgb1b:itgb1b-sGFP)* gill arches (left) and higher-magnification cross-sectional view of a lamella (right), highlighting Itgb1b-sGFP localization throughout the lamella, including the pavement cell basement membrane and pillar cells, with enrichment at the CA-CC interface. (C, D) Schematic of a pillar cell from Fig.2G (C) and close-up on an individual CA (red) surrounding a CC (white). (E) Detailed schematic representation of the regions highlighted in D: (1) CA–CC adhesion site within the pillar cell and (2) the autocellular tight junction sealing the pillar cell.

**Fig. S2.**
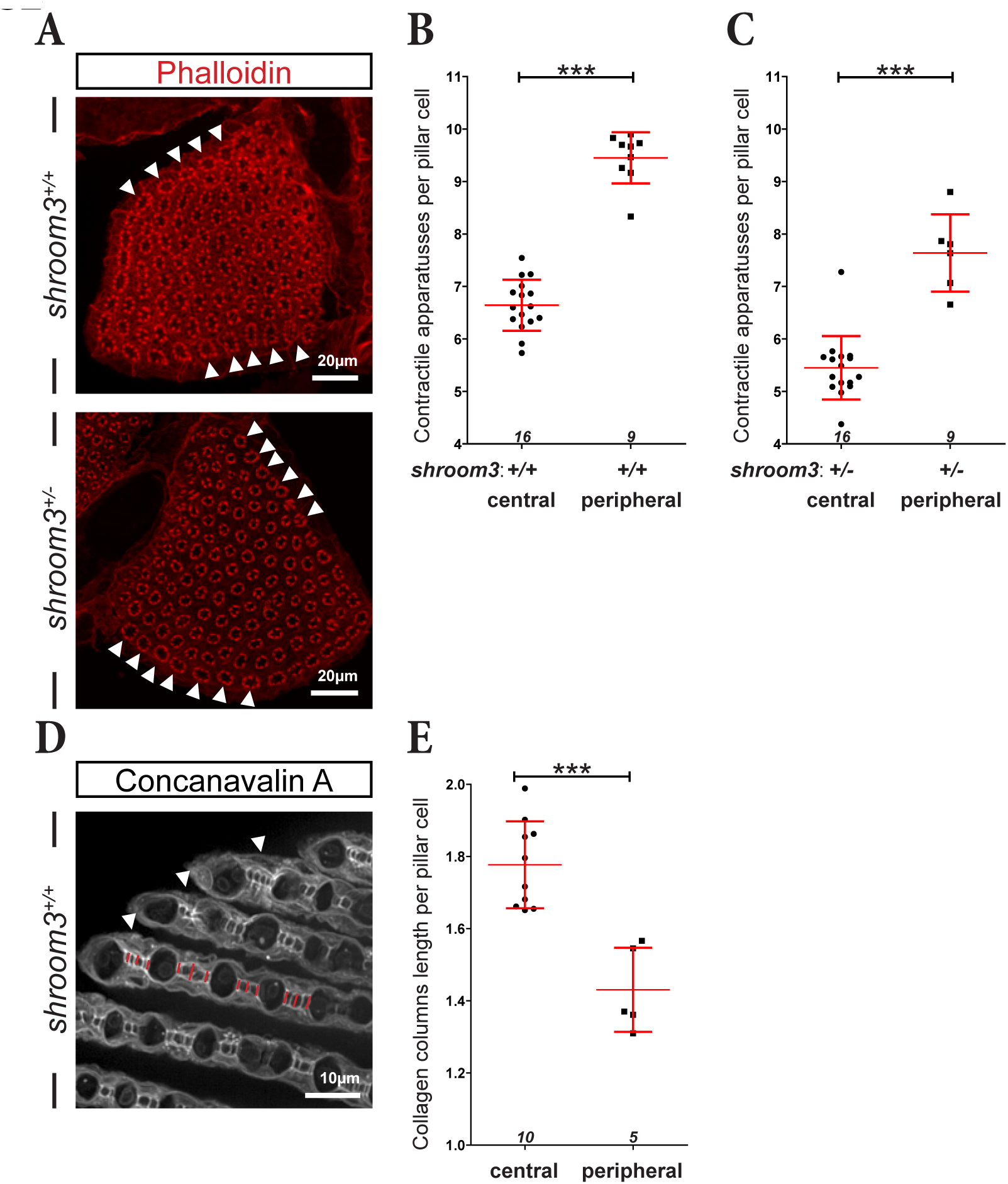
The pillar cell spatial pattern within the lamellae is related to their degree of constriction. (A) Representative MIPs of cryosections from adult gill arches labeled with phalloidin to visualize F-actin in *shroom3^+/+^* and *shroom3^+/-^* fish. Peripheral PCs (arrowheads) are located near the OMC, whereas central PCs are located within the lamella. (B, C) Quantification of CA number per central and peripheral PCs in *shroom3^+/+^* (B) and *shroom3^+/-^* (C). Values of central PC are taken from Fig. 3B. (D) Representative MIP of ECi-cleared gill cross-section from a *shroom3^+/+^* fish stained with Concanavalin A to label CCs. Peripheral PCs are marked by arrowheads at the distal end of the lamellae. Red lines indicate the CC lengths that were quantified. (E) Quantification of CC length in central and peripheral PCs in *shroom3^+/+^* fish. Values for central PC are taken from Fig. 3F. (B, C, E) Error bars indicate mean ± SD. Statistical significance was assessed using a two-tailed Mann-Whitney U test: \*\*\**P* ≤ 0.001. Each data point represents mean value of 90 PCs from three images of one fish. Numbers at the bottom of each panel indicate the number of animals analyzed (n).

**Fig. S3.**
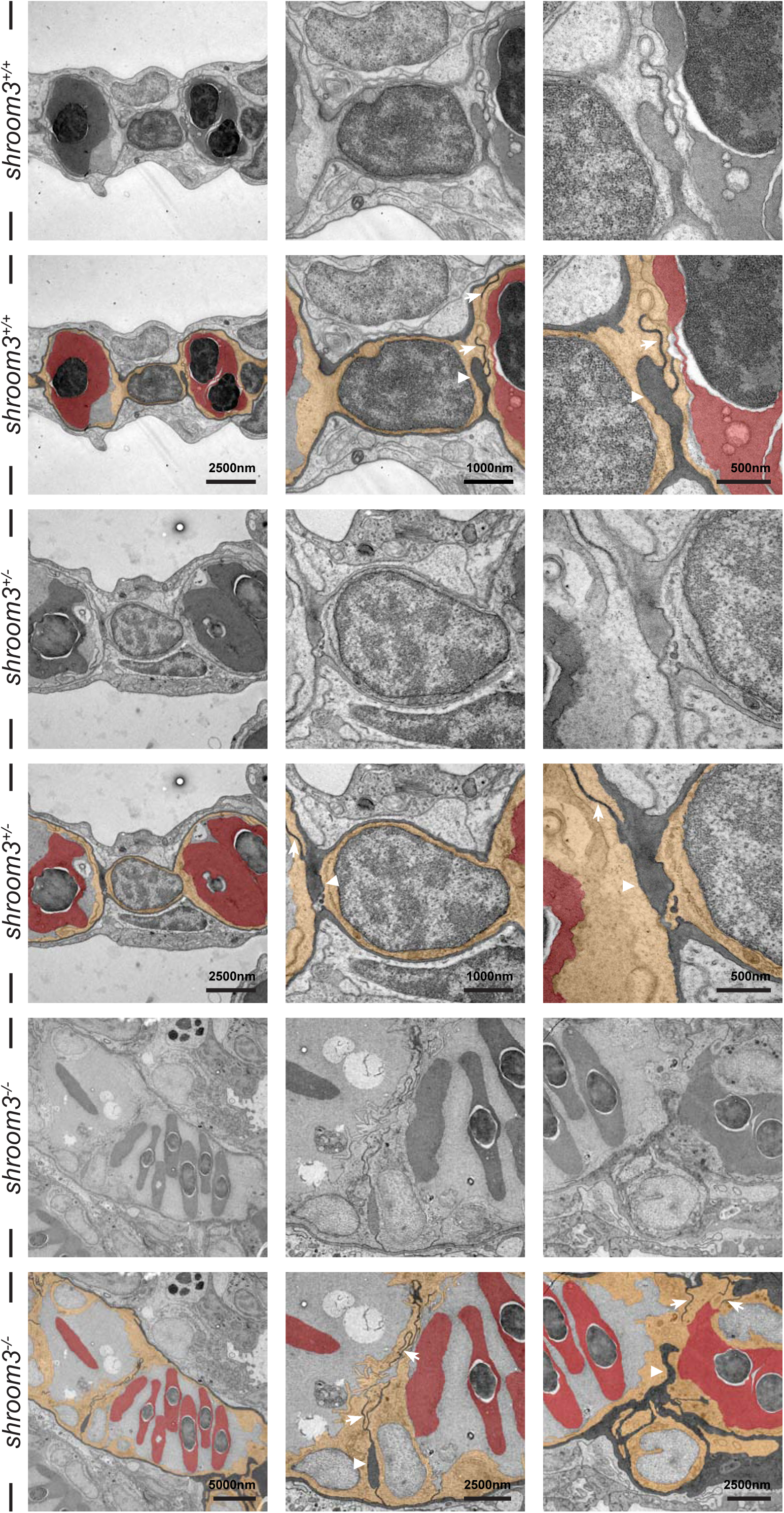
Ultrastructure of pillar cells across *shroom3* genotypes. Transmission electron micrographs of lamellar cross-sections from the indicated *shroom3* genotypes. For each genotype, the upper row shows the original micrographs, progressing from an overview of PC organization within the lamella (left) to higher-magnification views of PCs (middle) and detailed views of CCs (right). The lower row shows the same corresponding pseudocolored images, highlighting erythrocytes in red, PCs in yellow and the ECM in black. Arrowheads point on CCs and arrows on tight junctions.

**Fig. S4.**
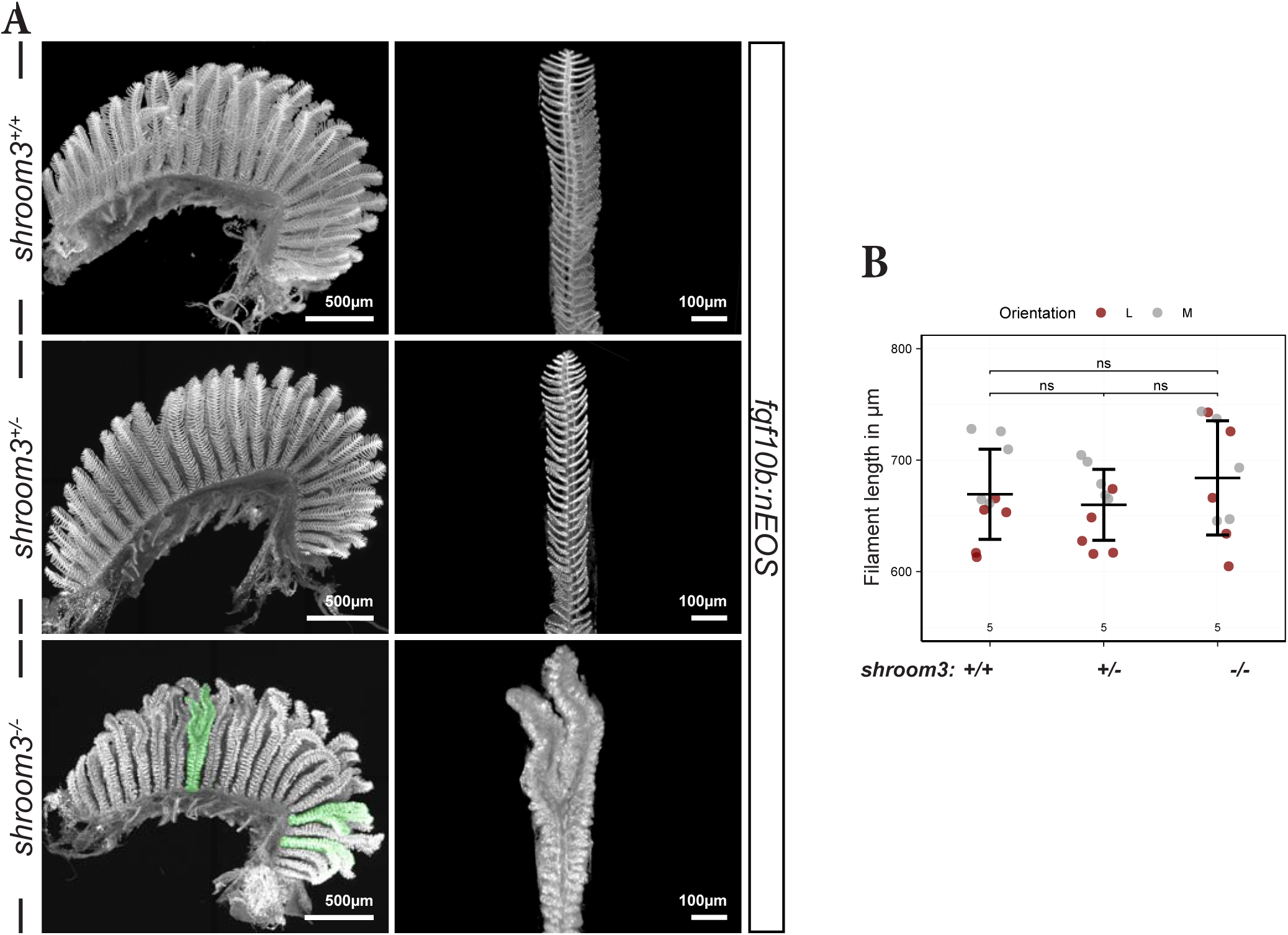
Loss of *shroom3* results in distal branching of gill filaments. (A) Representative MIPs of cleared *fgf10b:nEOS* first gill arches (left panels) and digitally isolated filaments (right) across all *shroom3* genotypes. Branched filaments in *shroom3^-/-^*mutants are highlighted in green in the whole arch and in the corresponding isolated filaments. (B) Length of lateral (red) and medial (white) filaments in the first gill arch of 6-month-old zebrafish across the indicated *shroom3* genotypes. Error bars indicate mean ± SD. Statistical significance was assessed using a two-tailed Mann-Whitney U test: ns, not significant. Numbers at the bottom of each panel indicate the number of animals analyzed (n).

**Fig. S5.**
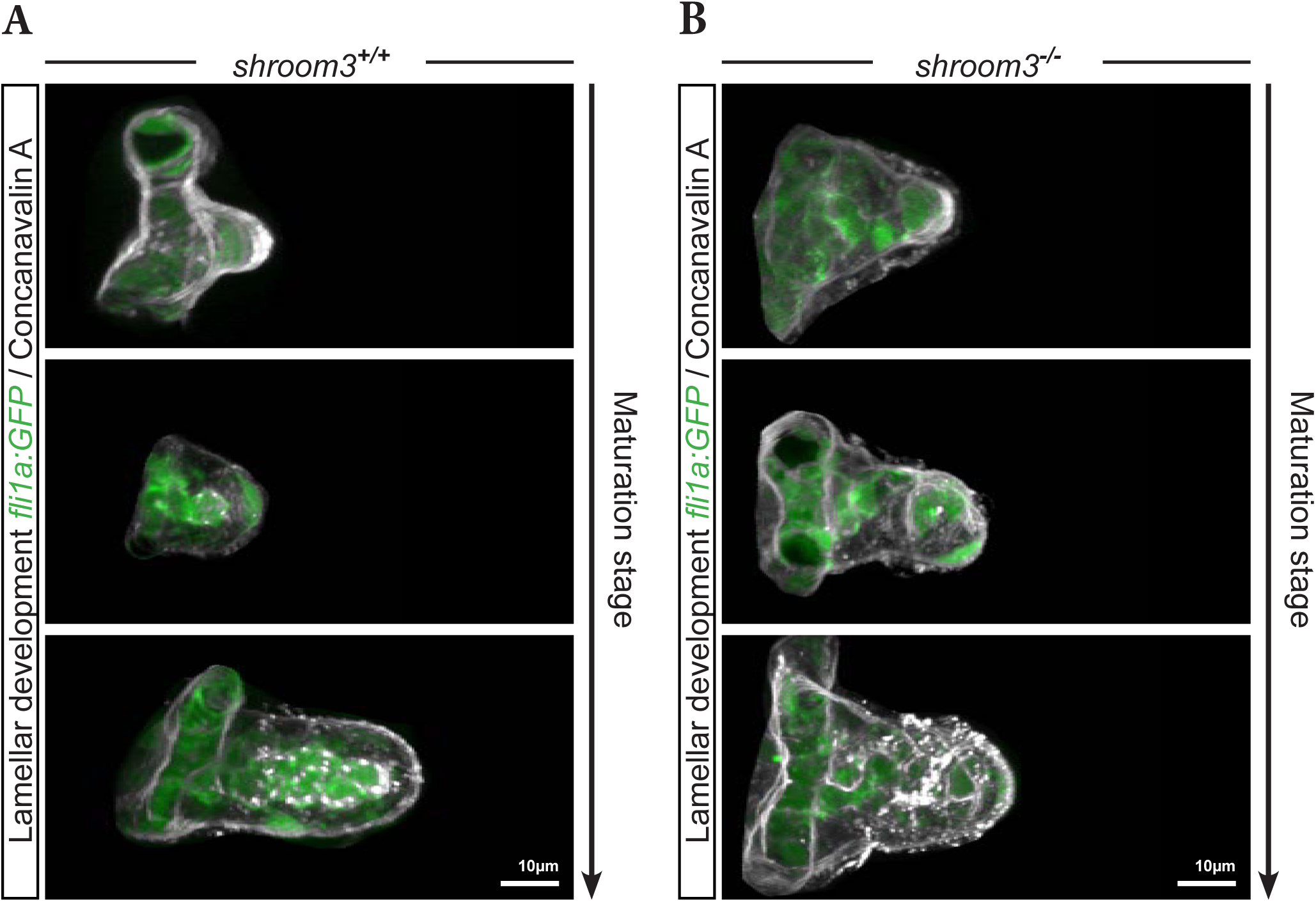
Collagen columns arise simultaneously with pillar cells during lamellar outgrowth. (A-B) Representative MIPs of lamellae at different stages of maturation in 14-dpf *fli1a:GFP shroom3^+/+^* (A) and *shroom3^-/-^* (B) larvae labeled with Concanavalin A. The first visible PC (*flia1a:GFP* positive) has already formed visible CCs, marked by Concanavalin A.

**Fig. S6.**
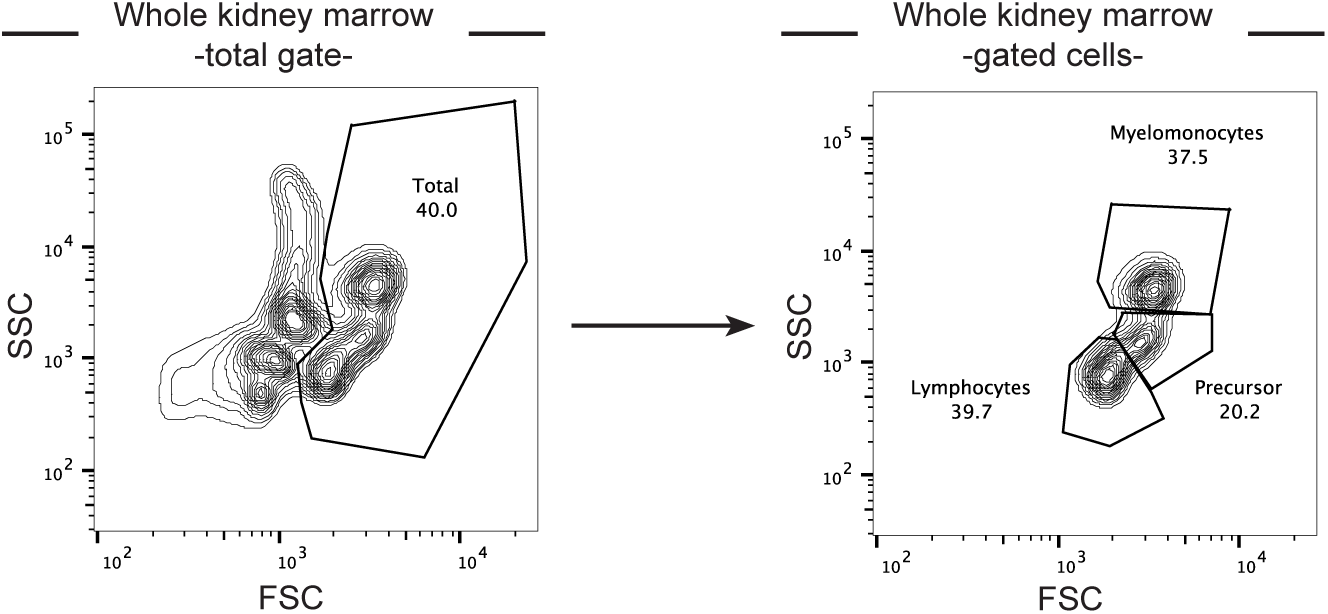
Whole kidney marrow gating strategy. Representative contour plots illustrating the gating strategy for WKM cells. Total cells were defined by forward scatter (FSC) and side scatter (SSC) while excluding erythrocytes and cellular debris (left plot). Within the total cell gate, myelomonocytes, precursor cells, and lymphocytes were identified based on their characteristic FSC and SSC profiles (right plot).

## Movie legends

**Movie 1: Digital isolation of an individual *shroom3^+/+^* lamella and subsequent quantification of pillar cell number and distribution**

Representative workflow demonstrating the digital segmentation of a single lamella from cleared *fgf10b:nEOS* gills and subsequent quantification of pillar cell number and spatial distribution in dorsal medial filaments of *shroom3^+/+^* fish (related to Fig. 4B).

**Movie 2: Digital isolation of an individual *shroom3^-/-^* lamella and subsequent quantification of pillar cell number and distribution**

Representative workflow demonstrating the digital segmentation of a single lamella from cleared *fgf10b:nEOS* gills and subsequent quantification of pillar cell number and spatial distribution in dorsal medial filaments of shroom3^-/-^ fish (related to Fig. 4B).

---

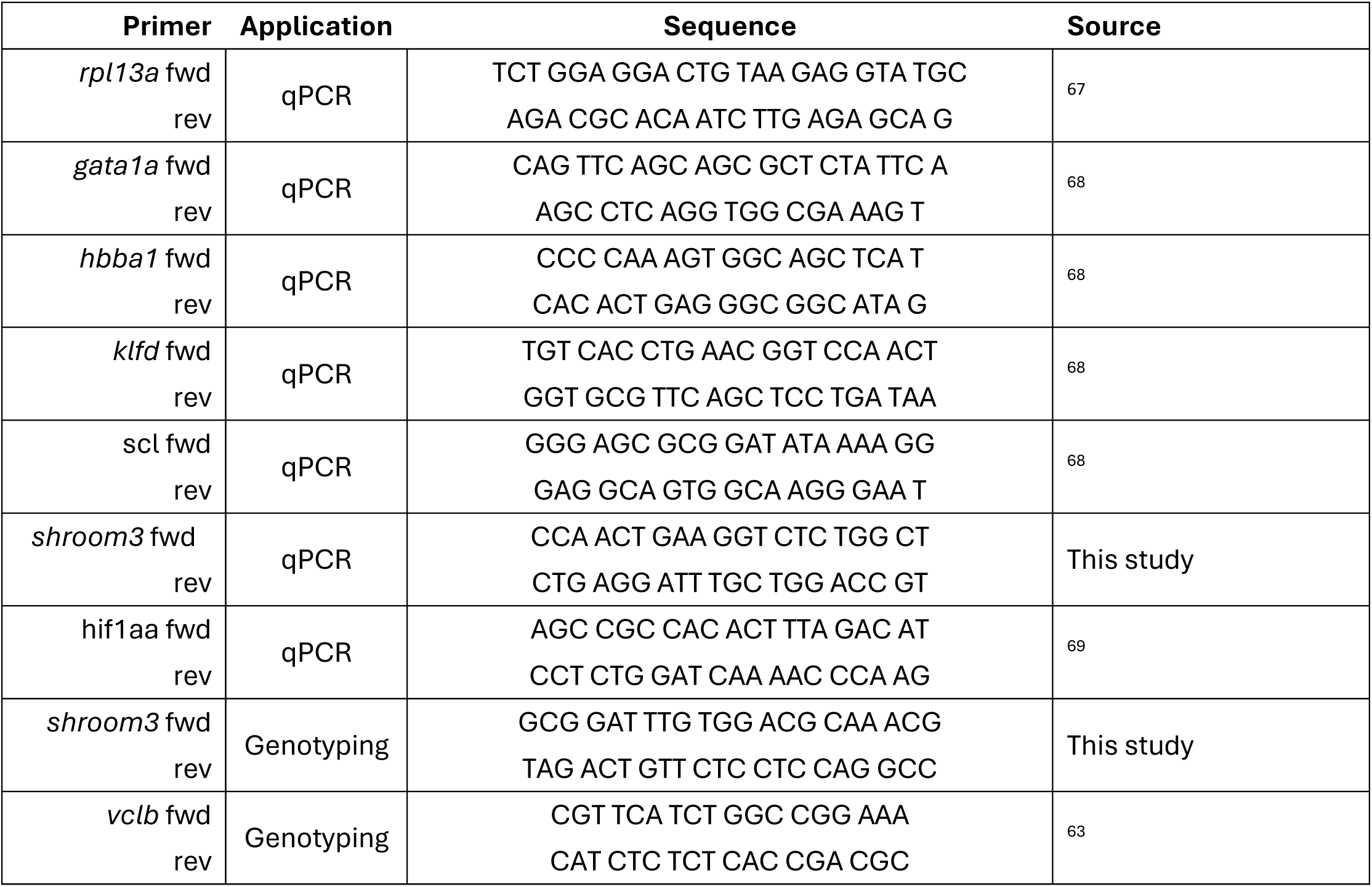

## References

1. Eckert, J., Viasnoff, V., and Yap, A.S. (2025). New directions in epithelial mechanoadaptation. Current Opinion in Cell Biology 95, 102536. 10.1016/j.ceb.2025.102536.

2. Kalukula, Y., Ciccone, G., Mohammed, D., Procès, A., Versaevel, M., Deridoux, A., Ergot, L., Barbier, Z., Mansy, M., Aucouturier, R., et al. (2025). Unlocking the therapeutic potential of cellular mechanobiology. Science Advances 11, eaea6817. 10.1126/sciadv.aea6817.

3. Panciera, T., Azzolin, L., Cordenonsi, M., and Piccolo, S. (2017). Mechanobiology of YAP and TAZ in physiology and disease. Nat Rev Mol Cell Biol 18, 758–770. 10.1038/nrm.2017.87.

4. Munjal, A., and Lecuit, T. (2014). Actomyosin networks and tissue morphogenesis. Development 141, 1789–1793. 10.1242/dev.091645.

5. Vicente-Manzanares, M., Ma, X., Adelstein, R.S., and Horwitz, A.R. (2009). Non-muscle myosin II takes centre stage in cell adhesion and migration. Nat Rev Mol Cell Biol 10, 778–790. 10.1038/nrm2786.

6. Costa, M., Wilson, E.T., and Wieschaus, E. (1994). A putative cell signal encoded by the *folded gastrulation* gene coordinates cell shape changes during Drosophila gastrulation. Cell 76, 1075–1089. 10.1016/0092-8674(94)90384-0.

7. Hildebrand, J.D., and Soriano, P. (1999). Shroom, a PDZ domain-containing actin-binding protein, is required for neural tube morphogenesis in mice. Cell 99, 485–497. 10.1016/s0092-8674(00)81537-8.

8. Lecuit, T., and Lenne, P.-F. (2007). Cell surface mechanics and the control of cell shape, tissue patterns and morphogenesis. Nat Rev Mol Cell Biol 8, 633–644. 10.1038/nrm2222.

9. Martin, A.C., and Goldstein, B. (2014). Apical constriction: themes and variations on a cellular mechanism driving morphogenesis. Development 141, 1987–1998. 10.1242/dev.102228.

10. Miao, H., and Blankenship, J.T. (2020). The pulse of morphogenesis: actomyosin dynamics and regulation in epithelia. Development 147, dev186502. 10.1242/dev.186502.

11. Suzuki, M., Morita, H., and Ueno, N. (2012). Molecular mechanisms of cell shape changes that contribute to vertebrate neural tube closure. Dev Growth Differ 54, 266–276. 10.1111/j.1440-169X.2012.01346.x.

12. Chung, M.-I., Nascone-Yoder, N.M., Grover, S.A., Drysdale, T.A., and Wallingford, J.B. (2010). Direct activation of Shroom3 transcription by Pitx proteins drives epithelial morphogenesis in the developing gut. Development 137, 1339–1349. 10.1242/dev.044610.

13. Plageman, T.F., Chauhan, B.K., Yang, C., Jaudon, F., Shang, X., Zheng, Y., Lou, M., Debant, A., Hildebrand, J.D., and Lang, R.A. (2011). A Trio-RhoA-Shroom3 pathway is required for apical constriction and epithelial invagination. Development 138, 5177–5188. 10.1242/dev.067868.

14. Ernst, S., Liu, K., Agarwala, S., Moratscheck, N., Avci, M.E., Dalle Nogare, D., Chitnis, A.B., Ronneberger, O., and Lecaudey, V. (2012). Shroom3 is required downstream of FGF signalling to mediate proneuromast assembly in zebrafish. Development 139, 4571–4581. 10.1242/dev.083253.

15. Khalili, H., Sull, A., Sarin, S., Boivin, F.J., Halabi, R., Svajger, B., Li, A., Cui, V.W., Drysdale, T., and Bridgewater, D. (2016). Developmental Origins for Kidney Disease Due to Shroom3 Deficiency. JASN 27, 2965–2973. 10.1681/ASN.2015060621.

16. Köttgen, A., Pattaro, C., Böger, C.A., Fuchsberger, C., Olden, M., Glazer, N.L., Parsa, A., Gao, X., Yang, Q., Smith, A.V., et al. (2010). Multiple New Loci Associated with Kidney Function and Chronic Kidney Disease: The CKDGen consortium. Nat Genet 42, 376–384. 10.1038/ng.568.

17. Matsuura, R., Hiraishi, A., Holzman, L.B., Hanayama, H., Harano, K., Nakamura, E., Hamasaki, Y., Doi, K., Nangaku, M., and Noiri, E. (2020). SHROOM3, the gene associated with chronic kidney disease, affects the podocyte structure. Sci Rep 10, 21103. 10.1038/s41598-020-77952-9.

18. Yeo, N.C., O’Meara, C.C., Bonomo, J.A., Veth, K.N., Tomar, R., Flister, M.J., Drummond, I.A., Bowden, D.W., Freedman, B.I., Lazar, J., et al. (2015). Shroom3 contributes to the maintenance of the glomerular filtration barrier integrity. Genome Res. 25, 57–65. 10.1101/gr.182881.114.

19. Kassianidou, E., Hughes, J.H., and Kumar, S. (2017). Activation of ROCK and MLCK tunes regional stress fiber formation and mechanics via preferential myosin light chain phosphorylation. MBoC 28, 3832–3843. 10.1091/mbc.e17-06-0401.

20. Nishimura, T., and Takeichi, M. (2008). Shroom3-mediated recruitment of Rho kinases to the apical cell junctions regulates epithelial and neuroepithelial planar remodeling. Development 135, 1493–1502. 10.1242/dev.019646.

21. Kato, A., Nakamura, K., Kudo, H., Tran, Y.H., Yamamoto, Y., Doi, H., and Hirose, S. (2007). Characterization of the Column and Autocellular Junctions That Define the Vasculature of Gill Lamellae. J Histochem Cytochem 55, 941–953. 10.1369/jhc.6A7154.2007.

22. Bettex-Galland, M., and Hughes, G.M. (1973). Contractile filamentous material in the pillar cells of fish gills. J Cell Sci 13, 359–370. 10.1242/jcs.13.2.359.

23. Kudo, H., Kato, A., and Hirose, S. (2007). Fluorescence visualization of branchial collagen columns embraced by pillar cells. J Histochem Cytochem 55, 57–62. 10.1369/jhc.6A7047.2006.

24. Karlsson, L. (1983). Gill morphology in the zebrafish, Brachydanio rerio (Hamilton- Buchanan). Journal of Fish Biology 23, 511–524. 10.1111/j.1095-8649.1983.tb02931.x.

25. Fabian, P., Tseng, K.-C., Thiruppathy, M., Arata, C., Chen, H.-J., Smeeton, J., Nelson, N., and Crump, J.G. (2022). Lifelong single-cell profiling of cranial neural crest diversification in zebrafish. Nat Commun 13, 13. 10.1038/s41467-021-27594-w.

26. Mongera, A., Singh, A.P., Levesque, M.P., Chen, Y.-Y., Konstantinidis, P., and Nüsslein-Volhard, C. (2013). Genetic lineage labeling in zebrafish uncovers novel neural crest contributions to the head, including gill pillar cells. Development 140, 916–925. 10.1242/dev.091066.

27. Park, J.S., Gutkowski, J., Castranova, D., Kenton, M., Sur, A., Martinez-Aceves, C., Nguyen, M.-A., Dell, C., Margolin, G., Iben, J., et al. (2025). Specialized gas-exchange endothelium of the zebrafish gill. Preprint at bioRxiv, 10.64898/2025.11.30.690480 https://doi.org/10.64898/2025.11.30.690480.

28. Yamaguchi, N., Zhang, Z., Schneider, T., Wang, B., Panozzo, D., and Knaut, H. (2022). Rear traction forces drive adherent tissue migration in vivo. Nat Cell Biol 24, 194–204. 10.1038/s41556-022-00844-9.

29. Stensløkken, K.-O., Sundin, L., and Nilsson, G.E. (2006). Endothelin receptors in teleost fishes: cardiovascular effects and branchial distribution. Am J Physiol Regul Integr Comp Physiol 290, 852–860. 10.1152/ajpregu.00618.2004.

30. Preußner, M., Mertens, A., Basoglu, M., and Lecaudey, V. (2025). A developmental atlas of zebrafish gills links early vascular patterning to adult architecture. Development 152, dev204984. 10.1242/dev.204984.

31. Shi, L., Chen, L., Jiang, S., Wu, Z., Zhou, Y., and Xu, Q. (2022). Hematogenesis Adaptation to Long-Term Hypoxia Acclimation in Zebrafish (Danio rerio). Fishes 7, 98. 10.3390/fishes7030098.

32. Traver, D., Paw, B.H., Poss, K.D., Penberthy, W.T., Lin, S., and Zon, L.I. (2003). Transplantation and in vivo imaging of multilineage engraftment in zebrafish bloodless mutants. Nat Immunol 4, 1238–1246. 10.1038/ni1007.

33. Haigo, S.L., Hildebrand, J.D., Harland, R.M., and Wallingford, J.B. (2003). Shroom Induces Apical Constriction and Is Required for Hingepoint Formation during Neural Tube Closure. Current Biology 13, 2125–2137. 10.1016/j.cub.2003.11.054.

34. Hildebrand, J.D. (2005). Shroom regulates epithelial cell shape via the apical positioning of an actomyosin network. Journal of Cell Science 118, 5191–5203. 10.1242/jcs.02626.

35. Liu, X., Liu, J., Wang, Y., Yao, M., Baker, K.B., Klapholz, B., Brown, N.H., Goult, B.T., and Yan, J. (2025). The mechanical response of vinculin. Science Advances 11, eady6949. 10.1126/sciadv.ady6949.

36. Gillich, A., Zhang, F., Farmer, C.G., Travaglini, K.J., Tan, S.Y., Gu, M., Zhou, B., Feinstein, J.A., Krasnow, M.A., and Metzger, R.J. (2020). Capillary cell-type specialization in the alveolus. Nature 586, 785–789. 10.1038/s41586-020-2822-7.

37. Mitra, S.K., Hanson, D.A., and Schlaepfer, D.D. (2005). Focal adhesion kinase: in command and control of cell motility. Nat Rev Mol Cell Biol 6, 56–68. 10.1038/nrm1549.

38. Morgan, M. (1974). Development of secondary lamellae of the gills of the trout, Salmo gairdneri (Richardson). Cell Tissue Res. 151. 10.1007/BF00222996.

39. Stensløkken, K.-O., Sundin, L., and Nilsson, G.E. (1999). Cardiovascular and gill microcirculatory effects of endothelin-1 in atlantic cod: evidence for pillar cell contraction. J Exp Biol 202, 1151–1157. 10.1242/jeb.202.9.1151.

40. Stolper, J., Ambrosio, E.M., Danciu, D.-P., Buono, L., Elliott, D.A., Naruse, K., Martínez-Morales, J.R., Marciniak-Czochra, A., and Centanin, L. (2019). Stem cell topography splits growth and homeostatic functions in the fish gill. eLife 8, e43747. 10.7554/eLife.43747.

41. McGreevy, E.M., Vijayraghavan, D., Davidson, L.A., and Hildebrand, J.D. (2015). Shroom3 functions downstream of planar cell polarity to regulate myosin II distribution and cellular organization during neural tube closure. Biol Open 4, 186–196. 10.1242/bio.20149589.

42. Carleton, J.L., Halabi, R.R., Willson, J.A., Jr, T.F.P., Bridgewater, D., Feng, Q., and Drysdale, T.A. (2025). Understanding the role of Shroom3 in the developing mouse myocardium. PLOS ONE 20, e0331583. 10.1371/journal.pone.0331583.

43. Durbin, M.D., O’Kane, J., Lorentz, S., Firulli, A.B., and Ware, S.M. (2020). SHROOM3 is downstream of the planar cell polarity pathway and loss-of-function results in congenital heart defects. Dev Biol 464, 124–136. 10.1016/j.ydbio.2020.05.013.

44. Newstead, J.D. (1967). Fine structure of the respiratory lamellae of teleostean gills. Z Zellforsch Mikrosk Anat 79, 396–428. 10.1007/BF00335484.

45. Sultana, N., Nag, K., Kato, A., and Hirose, S. (2007). Pillar cell and erythrocyte localization of fugu ETA receptor and its implication. Biochemical and Biophysical Research Communications 355, 149–155. 10.1016/j.bbrc.2007.01.128.

46. Liu, W., Xiu, L., Zhou, M., Li, T., Jiang, N., Wan, Y., Qiu, C., Li, J., Hu, W., Zhang, W., et al. (2024). The Critical Role of the Shroom Family Proteins in Morphogenesis, Organogenesis and Disease. Phenomics 4, 187–202. 10.1007/s43657-023-00119-9.

47. Kirschstein, T., Sahre, T., Kernig, K., Protzel, C., Porath, K., Köhling, R., and Hakenberg, O.W. (2015). Inverse relationship of Rho kinase and myosin-light chain kinase expression in the aging human detrusor smooth muscle. BMC Urol 15, 104. 10.1186/s12894-015-0098-2.

48. Totsukawa, G., Wu, Y., Sasaki, Y., Hartshorne, D.J., Yamakita, Y., Yamashiro, S., and Matsumura, F. (2004). Distinct roles of MLCK and ROCK in the regulation of membrane protrusions and focal adhesion dynamics during cell migration of fibroblasts. J Cell Biol 164, 427–439. 10.1083/jcb.200306172.

49. Totsukawa, G., Yamakita, Y., Yamashiro, S., Hartshorne, D.J., Sasaki, Y., and Matsumura, F. (2000). Distinct Roles of Rock (Rho-Kinase) and Mlck in Spatial Regulation of Mlc Phosphorylation for Assembly of Stress Fibers and Focal Adhesions in 3t3 Fibroblasts. J Cell Biol 150, 797–806. 10.1083/jcb.150.4.797.

50. Xu, L.-N., Sun, Y.-Y., Tan, Y.-F., Zhou, X.-Y., Xiang, T.-C., Fang, Y., Li, F., Shen, Q., Xu, H., and Rao, J. (2025). SHROOM3 Deficiency Aggravates Adriamycin-Induced Nephropathy Accompanied by Focal Adhesion Disassembly and Stress Fiber Disorganization. Cells 14, 895. 10.3390/cells14120895.

51. Atherton, P., Stutchbury, B., Jethwa, D., and Ballestrem, C. (2016). Mechanosensitive components of integrin adhesions: Role of vinculin. Exp Cell Res 343, 21–27. 10.1016/j.yexcr.2015.11.017.

52. Ezzell, R.M., Goldmann, W.H., Wang, N., Parashurama, N., Ingber, D.E., and New Collective Author (1997). Vinculin promotes cell spreading by mechanically coupling integrins to the cytoskeleton. Exp Cell Res 231, 14–26. 10.1006/excr.1996.3451.

53. Geiger, B., Tokuyasu, K.T., Dutton, A.H., and Singer, S.J. (1980). Vinculin, an intracellular protein localized at specialized sites where microfilament bundles terminate at cell membranes. Proceedings of the National Academy of Sciences 77, 4127–4131. 10.1073/pnas.77.7.4127.

54. Goldmann, W.H. (2016). Role of vinculin in cellular mechanotransduction. Cell Biol Int 40, 241–256. 10.1002/cbin.10563.

55. Johnson, R.P., and Craig, S.W. (1995). F-actin binding site masked by the intramolecular association of vinculin head and tail domains. Nature 373, 261–264. 10.1038/373261a0.

56. Le Clainche, C., Dwivedi, S.P., Didry, D., and Carlier, M.-F. (2010). Vinculin is a dually regulated actin filament barbed end-capping and side-binding protein. J Biol Chem 285, 23420–23432. 10.1074/jbc.M110.102830.

57. Wen, K.-K., Rubenstein, P.A., and DeMali, K.A. (2009). Vinculin Nucleates Actin Polymerization and Modifies Actin Filament Structure. J Biol Chem 284, 30463–30473. 10.1074/jbc.M109.021295.

58. Hadjaje, S., Andrade-Silva, I., Dalbe, M.-J., Clément, R., and Marthelot, J. (2024). Mechanics of Drosophila wing deployment. Nat Commun 15, 10577. 10.1038/s41467-024-54527-0.

59. Kimmel, C.B., Ballard, W.W., Kimmel, S.R., Ullmann, B., and Schilling, T.F. (1995). Stages of embryonic development of the zebrafish. Developmental Dynamics 203, 253–310. 10.1002/aja.1002030302.

60. Chi, N.C., Shaw, R.M., De Val, S., Kang, G., Jan, L.Y., Black, B.L., and Stainier, D.Y.R. (2008). Foxn4 directly regulates tbx2b expression and atrioventricular canal formation. Genes Dev 22, 734–739. 10.1101/gad.1629408.

61. Lawson, N.D., and Weinstein, B.M. (2002). *In Vivo* Imaging of Embryonic Vascular Development Using Transgenic Zebrafish. Developmental Biology 248, 307–318. 10.1006/dbio.2002.0711.

62. Maître, J.-L., Berthoumieux, H., Krens, S.F.G., Salbreux, G., Jülicher, F., Paluch, E., and Heisenberg, C.-P. (2012). Adhesion Functions in Cell Sorting by Mechanically Coupling the Cortices of Adhering Cells. Science 338, 253–256. 10.1126/science.1225399.

63. Gunawan, F., Gentile, A., Fukuda, R., Tsedeke, A.T., Jiménez-Amilburu, V., Ramadass, R., Iida, A., Sehara-Fujisawa, A., and Stainier, D.Y.R. (2019). Focal adhesions are essential to drive zebrafish heart valve morphogenesis. J Cell Biol 218, 1039–1054. 10.1083/jcb.201807175.

64. Lecaudey, V., Anselme, I., Rosa, F., and Schneider-Maunoury, S. (2004). The zebrafish Iroquois gene iro7 positions the r4/r5 boundary and controls neurogenesis in the rostral hindbrain. Development 131, 3121–3131. 10.1242/dev.01190.

65. Svec, D., Tichopad, A., Novosadova, V., Pfaffl, M.W., and Kubista, M. (2015). How good is a PCR efficiency estimate: Recommendations for precise and robust qPCR efficiency assessments. Biomol Detect Quantif 3, 9–16. 10.1016/j.bdq.2015.01.005.

66. Livak, K.J., and Schmittgen, T.D. (2001). Analysis of relative gene expression data using real-time quantitative PCR and the 2(-Delta Delta C(T)) Method. Methods 25, 402–408. 10.1006/meth.2001.1262.

67. Allanki, S., Strilic, B., Scheinberger, L., Onderwater, Y.L., Marks, A., Günther, S., Preussner, J., Kikhi, K., Looso, M., Stainier, D.Y.R., et al. Interleukin-11 signaling promotes cellular reprogramming and limits fibrotic scarring during tissue regeneration. Sci Adv 7, eabg6497. 10.1126/sciadv.abg6497.

68. Kulkeaw, K., Ishitani, T., Kanemaru, T., Fucharoen, S., and Sugiyama, D. (2010). Cold exposure down-regulates zebrafish hematopoiesis. Biochemical and biophysical research communications 394, 859–864. 10.1016/j.bbrc.2010.01.047.

69. Gerri, C., Marín-Juez, R., Marass, M., Marks, A., Maischein, H.-M., and Stainier, D.Y.R. (2017). Hif-1α regulates macrophage-endothelial interactions during blood vessel development in zebrafish. Nat Commun 8, 15492. 10.1038/ncomms15492.

